# OpenCell: proteome-scale endogenous tagging enables the cartography of human cellular organization

**DOI:** 10.1101/2021.03.29.437450

**Authors:** Nathan H. Cho, Keith C. Cheveralls, Andreas-David Brunner, Kibeom Kim, André C. Michaelis, Preethi Raghavan, Hirofumi Kobayashi, Laura Savy, Jason Y. Li, Hera Canaj, James Y.S. Kim, Edna M. Stewart, Christian Gnann, Frank McCarthy, Joana P. Cabrera, Rachel M. Brunetti, Bryant B. Chhun, Greg Dingle, Marco Y. Hein, Bo Huang, Shalin B. Mehta, Jonathan S. Weissman, Rafael Gómez-Sjöberg, Daniel N. Itzhak, Loic A. Royer, Matthias Mann, Manuel D. Leonetti

## Abstract

Elucidating the wiring diagram of the human cell is a central goal of the post-genomic era. We combined genome engineering, confocal live-cell imaging, mass spectrometry and data science to systematically map the localization and interactions of human proteins. Our approach provides a data-driven description of the molecular and spatial networks that organize the proteome. Unsupervised clustering of these networks delineates functional communities that facilitate biological discovery, and uncovers that RNA-binding proteins form a specific sub-group defined by unique interaction and localization properties. Furthermore, we discover that remarkably precise functional information can be derived from protein localization patterns, which often contain enough information to identify molecular interactions. Paired with a fully interactive website opencell.czbiohub.org, we provide a resource for the quantitative cartography of human cellular organization.

---

Sequencing the human genome has transformed cell biology by defining the protein parts list that forms the canvas of cellular operation (*1*, *2*). This paves the way for elucidating how the ~20,000 proteins encoded in the genome organize in space and time to define the cell’s functional architecture (*3*, *4*). Where does each protein localize within the cell? Can we comprehensively map how proteins assemble into larger functional communities? A main challenge to answering these fundamental questions is that cellular architecture is organized along multiple scales. Therefore, several approaches need to be combined for its elucidation (*5*). In a series of pioneering studies, human protein-protein interactions have been mapped using ectopic expression strategies with yeast two-hybrid (Y2H) (*6*) or epitope tagging coupled to immunoprecipitation-mass spectrometry (IP-MS) (*7*, *8*), while protein localization has been charted using immuno-fluorescence in fixed samples (*9*). A complementary approach is to directly modify genes in a genome by appending sequences that illuminate specific aspects of the corresponding proteins’ function (commonly referred to as “endogenous tagging” (*10*)). For example, endogenously tagging a gene with a fluorescent reporter enables to image protein sub-cellular localization in live cells, and supports functional characterization in a native cellular environment (*10*, *11*). The use of endogenous tagging to study the organization of a eukaryotic cell is illustrated by seminal work in the budding yeast *S. cerevisiae*. There, libraries of tagged strains have enabled the comprehensive mapping of protein localization and molecular interactions across the yeast proteome (*12*–*14*). These libraries were made possible by the relative simplicity of homologous recombination and genome engineering in yeast (*15*). In human cells, earlier work has leveraged alternative strategies including expression from bacterial artificial chromosomes (*16*) or central-dogma tagging (*17*) because of the difficulty of site-specific gene editing. CRISPR-mediated genome engineering now allows for homologous recombination-based endogenous tagging to be applied for the interrogation of the human cell (*10*, *11*, *18*).

Here, we combine experimental and analytical strategies to create OpenCell, a proteomic map of human cellular architecture. We generated a library of 1,310 CRISPR-edited HEK293T cell lines harboring fluorescent tags on individual proteins, which we characterized by pairing confocal microscopy and mass spectrometry. Our dataset constitutes the most comprehensive live-cell image collection of human protein localization to date. In addition, integration of IP-MS using the fluorescent tags for affinity capture enables measurement of localization and interactions from the same samples. For a quantitative description of cellular architecture, we introduce a data-driven framework to represent protein interactions and localization features, supported by a new machine learning algorithm for image encoding. This approach allows us to delineate communities of functionally related proteins by unsupervised clustering and facilitates the generation of mechanistic hypotheses, including for proteins that had so far remained uncharacterized. We further demonstrate that the localization pattern of each protein is defined by unique and specific features that can be used for functional interpretation, to the point that spatial relationships often contain enough information to predict interactions at the molecular scale. Finally, our analysis enables an unsupervised description of the human proteome’s organization, and highlights in particular that RNA-binding proteins exhibit unique functional signatures that shape the proteome’s network.

## Engineered cell library

Fluorescent protein (FP) fusions are versatile tools that can measure both protein localization by microscopy and protein-protein interactions by acting as affinity handles for IP-MS (*18*, *19*) (Fig. S1A). Here, we constructed a library of fluorescently tagged HEK293T cell lines by targeting human genes with the split-mNeonGreen2 system (*20*) (Fig. 1A). Split-FPs greatly simplify CRISPR-based genome engineering by circumventing the need for molecular cloning (*18*), and allowed us to generate endogenous genomic fusions (Fig. 1B) that preserve native expression regulation. A full description of our pipeline is available in the Methods section ((*21*); summarized in Fig. 1C through E). In brief, FP insertion sites (N- or C-terminus) were chosen on the basis of information from the literature or structural analysis (Fig. S1B; Table S1). For each tagged target we isolated a polyclonal pool of CRISPR-edited cells, which was then characterized by live-cell 3D confocal microscopy, IP-MS, and genotyping of tagged alleles by next-generation sequencing. Open-source software development and advances in instrumentation supported scalability (Fig. 1C). In particular, we developed *crispycrunch*, a CRISPR design software that enables guide RNA selection and homology donor sequence design (github.com/czbiohub/crispycrunch). We also fully automated the acquisition of data microscopy data in Python for on-the-fly computer vision and selection of desirable fields of view imaged in 96-well plates (github.com/czbiohub/2021-opencell-microscopy-automation). Our mass-spectrometry protocols use the high sensitivity of timsTOF instruments (*22*) which allowed miniaturization of IP-MS down to 0.8×10^6^ cells of starting material (Fig. S1C; about a tenth of the material required in previous approaches (*7*, *8*)).

**Figure 1:**
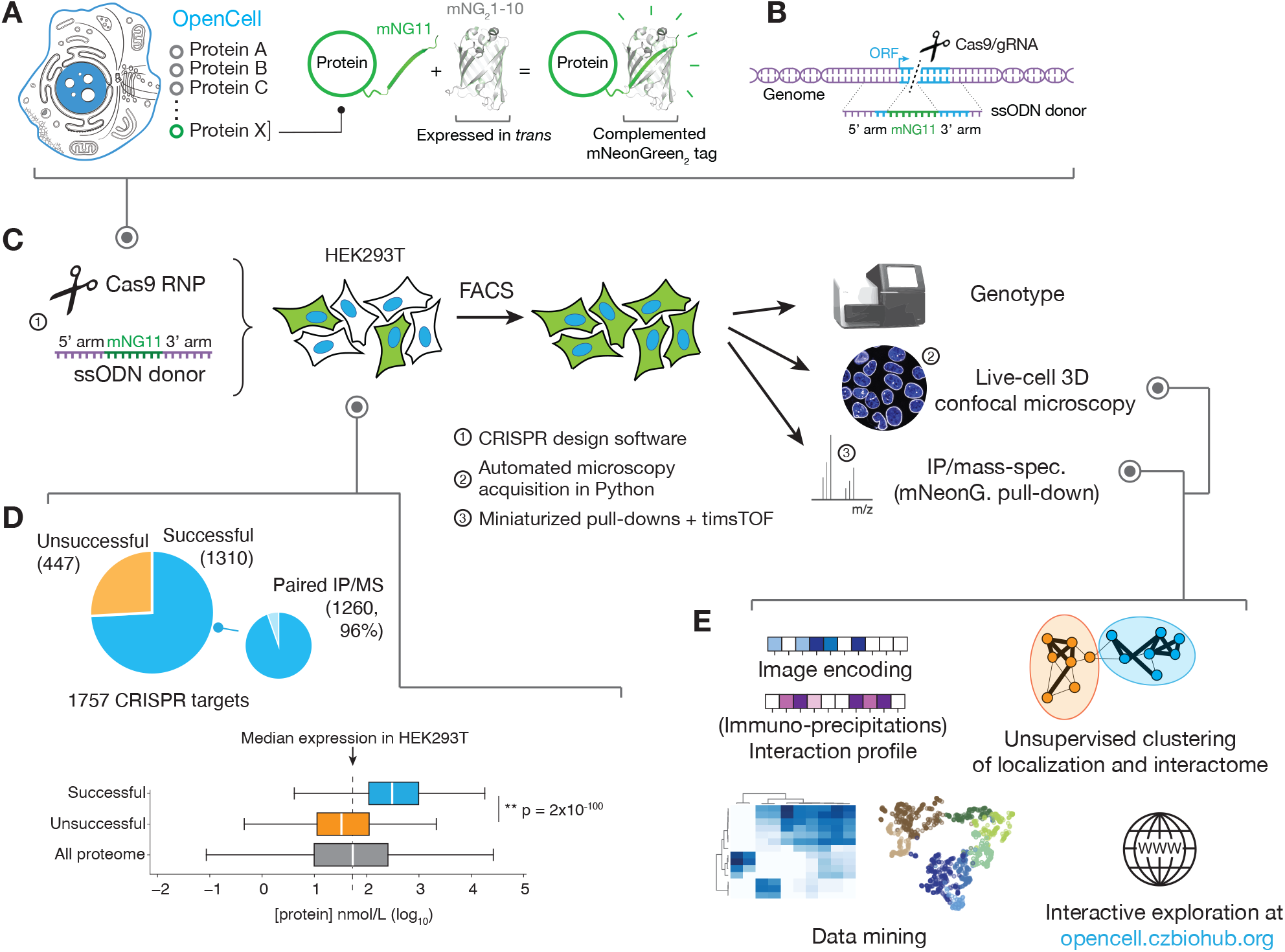
the OpenCell library. **(A)** Functional tagging with split-mNeonGreen_2_. In this system, mNeonGreen_2_ is separated into two fragments: a short mNG11 fragment, which is fused to a protein of interest, and a large mNG_2_1-10 fragment, which is expressed separately in trans (that is, tagging is done in cells that have been engineered to constitutively express mNG_2_1-10). **(B)** Endogenous tagging strategy: mNG11 fusion sequences are inserted directly within genomic open reading frames (ORFs) using CRISPR-Cas9 gene editing and homologous recombination with single-stranded oligonucleotides donors (ssODN). **(C)** The OpenCell experimental pipeline. See text for details. **(D)** Successful detection of fluorescence in the OpenCell library. Out of 1757 genes that were originally targeted, fluorescent signal was successfully detected for 1310 (top panel). Low protein abundance is the main obstacle to successful detection. Bottom left panel shows the full distribution of abundance for all proteins expressed in HEK293T vs. successfully or unsuccessfully detected OpenCell targets; boxes represent 25th, 50th, and 75th percentiles, and whiskers represent 1.5x interquartile range. Median is indicated by a white line. P-value: Student’s t-test. **(E)** The OpenCell data analysis pipeline, described in subsequent sections.

In total, we targeted 1757 genes, of which 1310 (75%) could be detected by fluorescence imaging and form our current dataset (full library details in Table S1). From these, we obtained paired IP-MS measurements for 1260 targets (96%, Fig. 1D). The 1310-protein collection includes a balanced representation of the pathways, compartments and functions of the human proteome (Fig. S1D), with the exception of processes specific to mitochondria, organellar lumen or extracellular matrix. Indeed, the split-FP system tags a gene of interest with a short sequence (mNG11) while a larger FP fragment (mNG_2_1-10) is expressed separately (Fig. 1A). In the version used here, the mNG_2_1-10 fragment is expressed in the nucleo-cytoplasm and prevents access to proteins inside organellar compartments. Membrane proteins can be tagged as long as one terminus extends in the nucleo-cytoplasm. In future iterations, other split systems that contain compartment-specific signal sequences could be used to target organellar lumen (*23*).

Fluorescent tagging was readily successful for essential genes, suggesting that FP fusions are well tolerated (Fig. S2A). To evaluate other factors contributing to successful fluorescent detection, we measured RNA and protein concentration in HEK293T cells (Fig. S2B; using a 24-fraction scheme for deep proteome quantification; see fully annotated proteome in Table S2). This revealed that protein abundance is the main limitation to detection (Fig. 1D, S2C; see details for unsuccessful targets in Table S3); most successful targets are among the top 50% most abundant (Fig. S2D). Gene-editing efficiency was another important factor: among well-expressed targets, failure was correlated with significantly lower rates of homologous recombination (Fig. S2E), which would impair the selection of edited cells by fluorescence-activated cell sorting (FACS). Training a regression model revealed that the combination of protein abundance and editing efficiency could predict successful detection with 82% accuracy.

To maximize throughput, we used a polyclonal strategy to select genome-edited cells by FACS. Polyclonal pools contain cells with distinct genotypes. HEK293T are pseudo-triploid (*24*) and a single edited allele is sufficient to confer fluorescence. Moreover, various DNA repair mechanisms compete with homologous recombination for the resolution of CRISPR-induced genomic breaks (*25*) so that alleles containing non-functional mutations can be present in addition to the desired fusion alleles. However, such alleles do not support fluorescence and are therefore unlikely to impact other measurements, especially in the context of a polyclonal pool. We developed a stringent selection scheme to significantly enrich for fluorescent fusion alleles (Fig. S3A). Our final cell library has a median 61% of mNeonGreen-integrated alleles, 5% wild-type and 26% other non-functional alleles (Fig. S3B, full genotype information in Table S1).

Finally, we verified that our engineering approach maintained the endogenous abundance of the tagged target proteins. For this, we quantified protein expression by Western blotting using antibodies specific to proteins targeted in 12 different cell pools (Fig. S3C), and by single-shot mass spectrometry in 63 tagged lines (Fig. S3D). Both approaches revealed a median abundance of tagged targets in engineered lines at about 80% of untagged HEK293T control, with 5 outliers (8% of total) identified by proteomics (Fig. S3D, all within 3.5-fold of control). Importantly, the overall proteome composition was unchanged in all tagged lines (Fig. S3E-F). Overall, our gene-editing strategy preserves near-endogenous abundances and circumvents the limitations of ectopic overexpression (*11*, *26*, *27*), which include aberrant localization, changes in organellar morphology, and masking effects (see the examples of SPTLC1, TOMM20 and MAP1LC3B in Fig. S3G). Therefore, OpenCell supports the functional profiling of tagged proteins in their native cellular context.

## Interactome analysis and stoichiometry-driven clustering

Affinity enrichment coupled to mass spectrometry is an efficient and sensitive method for the systematic mapping of protein interaction networks (*28*). We isolated tagged proteins (“baits”) from cell lysates solubilized in digitonin, a mild non-ionic detergent that preserves the native structure and properties of membrane proteins (*29*). Specific protein interactors (“preys”) were identified by proteomics from biological triplicate experiments (see Figure S4A-B and (*21*) for a detailed description of our statistical analysis, which builds upon established methods (*7*)). In total, the full interactome from our 1260 OpenCell baits includes 29,922 interactions between 5292 proteins (baits and preys, Fig. 2A, full interactome data in Table S4). To assess the quality of our interactome, we estimated its precision (the fraction of true positive interactions over all interactions) and recall (the fraction of interactions identified compared to a ground truth set) using reference data (Fig. S4B). For recall analysis, we quantified the coverage in our data of interactions included in CORUM (*30*), a compendium of protein interactions manually curated from the literature. To estimate precision, we quantified how many of our interactions involved protein pairs expected to localize to the same broad cellular compartment (*31*) (Fig. S4B). To benchmark OpenCell against other large-scale interactomes, we compared its precision and recall to Bioplex (over-expression of HA-tagged baits (*8*, *32*)), the yeast-two-hybrid human reference interactome (HuRI (*6*)) and our own previous data (GFP fusions expressed from bacterial artificial chromosomes (*7*)) (Fig. S4C-E). We also calculated compression rates for each dataset as a measure of the overall richness in network patterns and motifs distinguishable from noise, which correlates with overall network quality: real-world networks contain redundant information which can be compressed, while pure noise is not compressible (see (*33*)) (Fig. S4F). Across all metrics, OpenCell outperformed previous approaches. OpenCell also includes many interactions not reported in previous datasets (Fig. S4E,G). Our interactome may better reflect biological interactions because it preserves near-endogenous protein expression.

**Figure 2:**
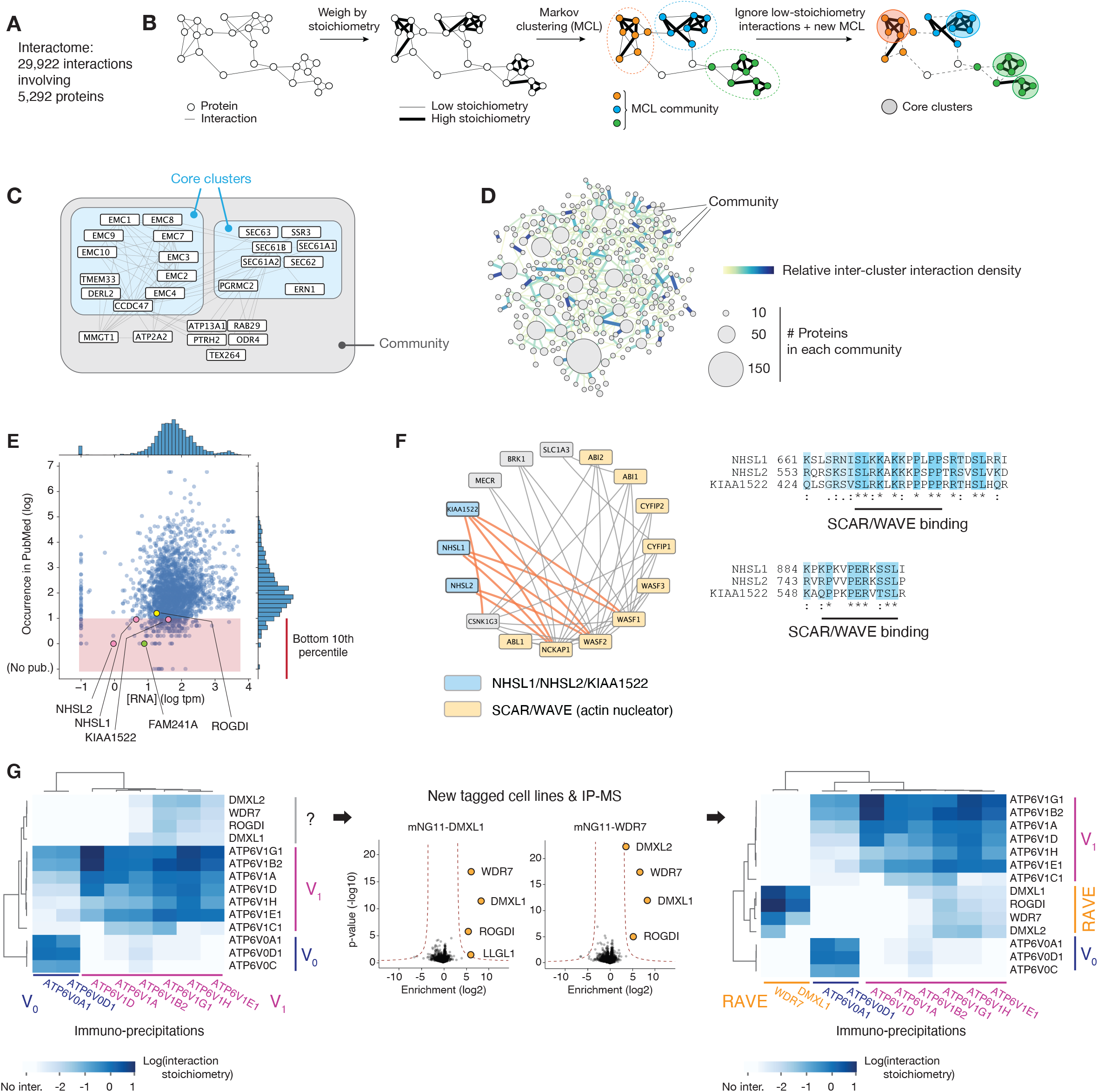
Protein interactome. **(A)** Overall description of the interactome. **(B)** Unsupervised Markov clustering of the interactome graph. **(C)** Example of community and core cluster definition for the translocon/EMC community. **(D)** The complete graph of connections between interactome communities. The density of protein-protein interactions between communities is represented by increased edge width. The numbers of targets included in each community is represented by circles of increasing diameters. **(E)** Distribution of occurrence in PubMed articles vs. RNA expression for all proteins found within interactome communities. The bottom 10th percentile of publication count (poorly characterized proteins) is highlighted. **(F)** NHSL1/NSHL2/KIAA1522 are part of the SCAR/WAVE community and share amino-acid sequence homology (right panel). **(G)** DMXL1/2, WDR7 and ROGDI form the human RAVE complex. Heatmaps represent the interaction stoichiometry of preys (lines) in the pull-downs of specific OpenCell targets (columns). See text for details.

A powerful way to interpret interactomes is to identify communities of interactors (*8*, *13*). To this end, we applied unsupervised Markov clustering (MCL) (*34*) to the graph of interactions defined by our data (5292 baits and preys). We first measured the stoichiometry of each interaction, using a quantitative approach we previously established (*7*). Interaction stoichiometry measures the abundance of a protein interactor relative to the abundance of the bait in a given immuno-precipitation sample. We have shown that stoichiometry can be interpreted as a proxy for interaction strength, and that interactions can be classified between core (i.e. high) and low stoichiometries (*7*). In our current data, both high- and low-stoichiometry interactions were significantly enriched for proteins pairs sharing gene ontology annotations (Fig. S4H). Using stoichiometry to assign weights to the edges in the interaction graph (Fig. 2B), a first round of MCL delineated inter-connected protein communities and led to better clustering performance than clustering based on connectivity alone (Fig. S4I). To better delineate stable complexes, we further refined each individual MCL community by additional clustering while removing low-stoichiometry interactions. The resulting sub-clusters outline core interactions within existing communities (Fig. 2B). Figure 2C illustrates how this unsupervised approach enables to delineate functionally related proteins: all subunits of the machinery responsible for the translocation of newly translated proteins at the ER membrane (SEC61/62/63) and of the EMC (ER Membrane Complex) are grouped within respective core interaction clusters, but both are part of the same larger MCL community. This mirrors the recently appreciated co-translational role of EMC for insertion of transmembrane domains at the ER (*35*). Additional proteins that have only recently been shown to act co-translationally are found clustering with translocon or EMC subunits, including ERN1 (IRE1) (*36*) and CCDC47 (*37*, *38*). Thus, clustering can facilitate mechanistic exploration by grouping proteins involved in related pathways. Overall, we identified 300 communities including a total of 2096 baits and preys (full details in Table S4). Ontology analysis revealed that these communities are significantly enriched for specific cellular functions, supporting their biological relevance (82% of all communities are significantly enriched for specific biological process or molecular function GO ontology terms; see Table S5 for complete analysis). A graph of interactions between communities reveals a richly interconnected network (Fig. 2D), the structure of which outlines the global architecture of the human interactome (discussed further below).

A direct application of interactome clustering is to help elucidate the cellular roles of the many human proteins that remain poorly characterized (*39*). We identified poorly characterized proteins by quantifying their occurrence in article titles and abstracts from PubMed (Fig. 2E). Empirically, we determined that proteins in the bottom 10^th^ percentile of publication count (corresponding to less than 10 publications) are very poorly annotated (Fig. 2E). This set encompasses a total of 251 proteins found in interaction communities for which our dataset offers potential mechanistic insights. For example, the proteins NHSL1, NHSL2 and KIAA1522 are all found as part of a community centered around SCAR/WAVE, a large multi-subunit complex nucleating actin polymerization (Fig. 2F). All three proteins share sequence homology and are homologous to NHS (Fig. S5A), a protein mutated in patients with Nance-Horan syndrome. NHS interacts with SCAR/WAVE components to coordinate actin remodeling (*40*). Thus, NHSL1, NHSL2 and KIAA1522 also act to regulate actin assembly. A recent mechanistic study supports this hypothesis: NHSL1 localizes at the cell’s leading edge and directly binds SCAR/WAVE to negatively regulate its activity, reducing F-actin content in lamellipodia and inhibiting cell migration (*41*). The authors identified NHSL1’s SCAR/WAVE binding sites, and we find these sequences to be conserved in NSHL2 and KIA1522 (Fig. 2F). Therefore, our data suggests that both NHSL2 and KIAA1522 are also direct SCAR/WAVE binders and possible modulators of the actin cytoskeleton.

Our data also sheds light on the function of ROGDI, whose variants cause Kohlschuetter-Toenz syndrome (a recessive developmental disease characterized by epilepsy and psychomotor regression (*42*)). ROGDI appears in the literature because of its association with disease, but no study, to our knowledge, specifically determines its molecular function. We first observed that ROGDI’s interaction pattern closely matched that of three other proteins in our dataset: DMXL1, DMXL2 and WDR7 (Fig. 2G). This set exhibited a specific interaction signature with the v-ATPase lysosomal proton pump. All four proteins interact with soluble v-ATPase subunits (ATP6-V1), but not its intra-membrane machinery (ATP6-V0). DMXL1 and WDR7 interact with V1 v-ATPase, and their depletion in cells compromises lysosomal re-acidification (*43*). Sequence analysis showed that DMXL1 or 2, WDR7 and ROGDI are homologous to proteins from yeast and Drosophila involved in the regulation of assembly of the soluble V1 subunits onto the V0 transmembrane ATPase core (*44*, *45*) (Fig. S5B). In yeast, Rav1 and Rav2 (homologous to DMXL1/2 and ROGDI, respectively) form the stoichiometric RAVE complex, a soluble chaperone that regulates v-ATPase assembly (*45*). To assess the existence of a human RAVE-like complex, we generated new tagged cell lines for DMXL1 and 2, WDR7, and ROGDI. Because of the low abundance of these proteins, the localization of DMXL2 and ROGDI were not detectable but pulldowns of DMXL1 and WDR7 confirmed a stoichiometric interaction between DMXL1 and 2, WDR7 and ROGDI (Fig. 2G, right panels). No direct interaction between DXML1 and DMXL2 was detected, suggesting that they might nucleate two separate sub-complexes. Therefore, our data reveals a human RAVE-like complex comprising DMXL1 or 2, WDR7 and ROGDI, which we propose acts as a chaperone for v-ATPase assembly based on its yeast homolog. Altogether, these results illustrate how our data can facilitate the generation of new mechanistic hypotheses by combining quantitative analysis and literature curation.

## Image dataset: localization annotation and self-supervised machine learning

A key advantage of our cell engineering approach is to enable the characterization of each tagged protein in live, unperturbed cells. To profile localization, we performed spinning-disk confocal fluorescence microscopy (63x 1.47NA objective) under environmental control (37°C, 5% CO_2_), and imaged the 3D distribution of proteins in consecutive z-slices. Microscopy acquisition was fully automated in Python to enable scalability (Fig. S6A-B). In particular, we trained a computer vision model to identify fields of view (FOVs) with homogeneous cell density on-the-fly, which reduced experimental variation between images. Our dataset contains a collection of 6375 3D stacks (5 different FOVs for each target) and includes paired imaging of nuclei with live-cell Hoechst 33342 staining.

We manually annotated localization patterns by assigning each protein to one or more of 15 separate cellular compartments such as the nucleolus, centrosome or Golgi apparatus (Fig. 3A). Because proteins often populate multiple compartments at steady-state (*9*), we graded annotations using a three-tier system: grade 3 identifies prominent localization compartment(s), grade 2 represents less pronounced localizations, and grade 1 annotates weak localization patterns nearing our limit of detection (see Fig. S7A for two representative examples, full annotations in Table S6). Ignoring grade 1 annotations which are inherently less precise, 55% of proteins in our library were detected in multiple locations consistent with known functional relationships. for example, clear connections were observed between secretory compartments (ER, Golgi, vesicles, plasma membrane), or between cytoskeleton and plasma membrane (Fig. S7B, Table S6)). Many proteins are found in both nucleus and cytoplasm (21% of our library), highlighting the importance of the nucleo-cytoplasmic import and export machinery in shaping global cellular function (*46*, *47*). Importantly, because our split-FP system does not enable the detection of proteins in the lumen of organelles, multi-localization involving translocation across an organellar membrane (which is rare but does happen for mitochondrial or peroxisomal proteins) cannot be detected in our data.

**Figure 3:**
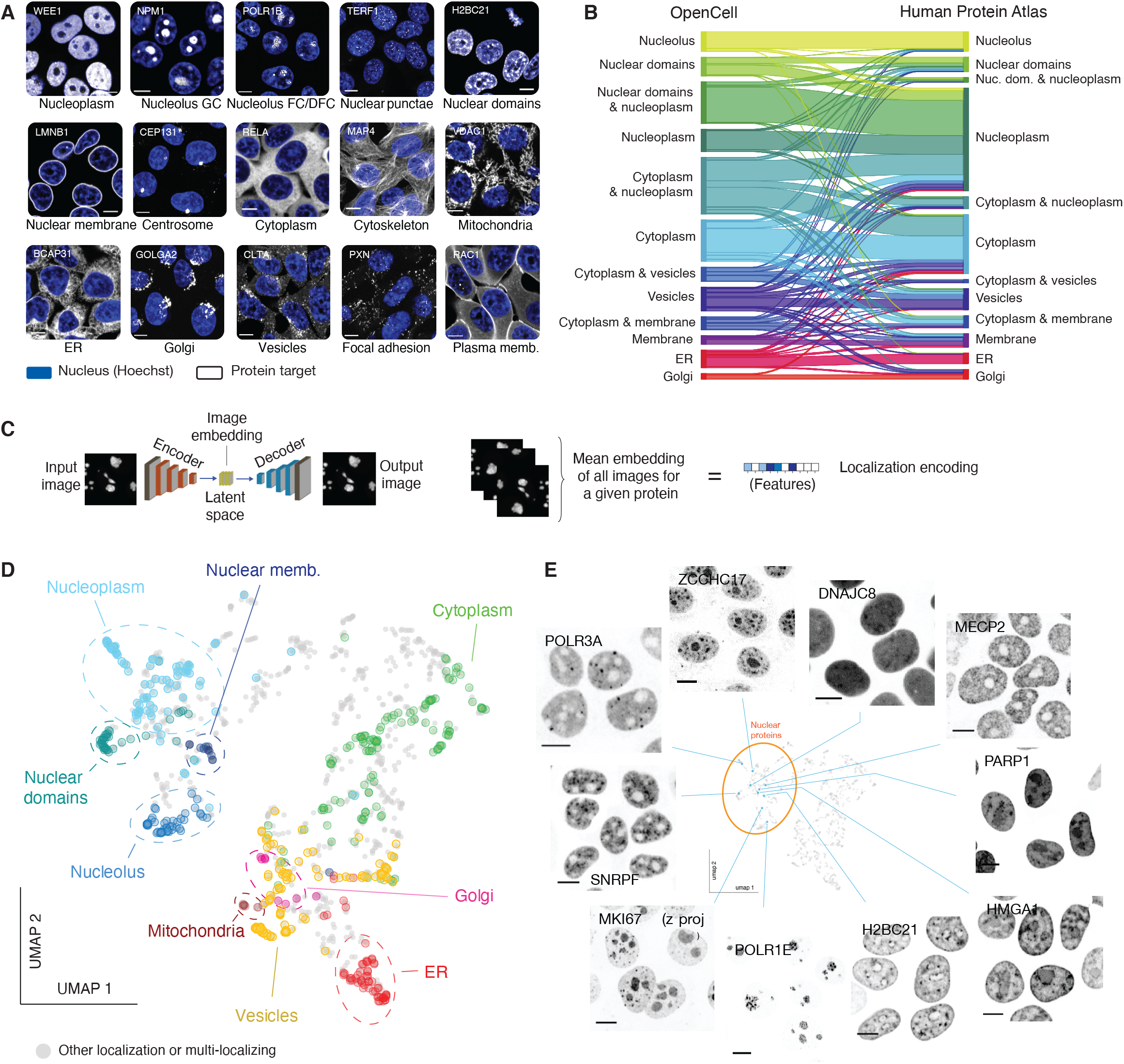
live-cell image collection. **(A)** The 15 cellular compartments segregated for annotating localization patterns. The localization of a representative protein belonging to each group is shown (greyscale, gene names in top left corners; scalebar: 10 μm). Nuclear stain (Hoechst) is shown in blue. “Nuclear domains” designate proteins with pronounced non-uniform nucleoplasmic localization, for example chromatin binding proteins. **(B)** Comparison of annotated localization for proteins included in both OpenCell and Human Protein Atlas datasets. In this flow diagram, colored bands represent groups of proteins that shared the same localization annotation in OpenCell, and the width of the band represents the number of proteins in each group. For readability, only the 12 most common localization groups are shown. Some multi-localization groups are included (e.g. “cytoplasm & nucleoplasm”). **(C)** Principle of localization encoding by self-supervised machine learning. See text for details. **(D)** UMAP representation of the OpenCell localization dataset, highlighting targets found to localize to a unique cellular compartment. **(E)** Representative images for 10 nuclear targets that exemplify the nuanced diversity of localization patterns across the proteome. Scale bars: 10 μm.

To benchmark our dataset, we compared our localization annotations against the Human Protein Atlas (HPA), the reference antibody-based compendium of human protein localization (*9*). This revealed significant agreement between datasets: 75% of proteins share at least one localization annotation in common (Fig. 3B; this includes 25% of all proteins that share the exact same set of annotations, see full description in Table S7A). Because HPA mostly reports on cell lines other than HEK293T, a perfect overlap is not expected as proteins might differentially localize between related compartments in different cell types. However, the annotations for 147 proteins (11% of our data) were fully inconsistent between the two datasets (Fig. S7C). An extensive curation of the literature on the localization of those proteins allowed us to resolve discrepancies for 115 proteins (i.e., 78% of that set; full curation in Table S8). Of these, existing literature evidence supported the OpenCell results for 113 (98.3%) of the 115 cases (Fig. S7D). This validates that endogenous tagging can help refine the curation of localization in the human proteome. Finally, our dataset includes 350 targets that have orthologs in *S. cerevisiae*. Comparison between OpenCell and yeast localization annotations (*48*) revealed a high degree of concordance (Fig. S7E; Table S7B; 81% of proteins share at least one annotation in common, including 36% perfect matches).

While expert annotation remains the best performing strategy to curate protein localization (*49*, *50*), the low-dimensional description it allows is not well suited for quantitative comparisons. Recent developments in image analysis and machine learning offer new opportunities to extract high-dimensional features from microscopy images (*50*, *51*). Therefore, we developed a deep learning model to quantitatively represent the localization pattern of each protein in our dataset (*52*). Briefly, our model is a variant of an autoencoder (Fig. 3C): a form of neural network that learns to vectorize an image through paired tasks of encoding (from an input image to a vector in a latent space) and decoding (from the latent space vector to a new output image). After training, a consensus representation for a given protein can be obtained from the average of the encodings from all its associated images. This generates a high-dimensional “localization encoding” (Fig. 3C) that captures the complex set of features that define the spatial distribution of a protein at steady state and across many individual cells. One of the main advantages of this approach is that it is self-supervised. Therefore, as opposed to supervised machine learning strategies that are trained to recognize pre-annotated patterns (for example, manual annotations of protein localization (*50*)), our method extracts localization signatures from raw images without any *a priori* assumptions or manually assigned labels. To visualize the relationships between these high-dimensional encodings, we embedded the encodings for all 1,310 OpenCell targets in two dimensions using UMAP, an algorithm that reduces high-dimensional datasets to two dimensions (UMAP 1 and UMAP 2) while attempting to preserve the global and local structures of the original data (*53*).The resulting map is organized in distinct territories that closely match manual annotations (Fig. 3D, highlighting mono-localizing proteins). This validates that the encoding approach yields a quantitative representation of the biologically relevant information in our microscopy data. The separation of different protein clusters in the UMAP embedding (further discussed below) mirrors the fascinating diversity of localization patterns across the full proteome. Images from nuclear proteins offer compelling illustrative examples of this diversity and reveal how fine-scale details can define the localization of proteins within the same organelle (Fig. 3E).

## Functional specificity of protein localization in the human cell

Extracting functional insights directly from cellular images is a major goal of modern cell biology and data science (*54*). In this context, our image library and associated machine learning encodings enable us to explore what degree of functional relationship can be inferred between proteins solely based on their localization. For this, we first employed an unsupervised Leiden clustering strategy commonly used to identify cell types in single-cell RNA sequencing datasets (*55*). Clusters group proteins that share similar localization properties (every protein in the dataset is included in a cluster); these groups can then be analyzed for how well they match different sets of groundtruth annotations (Fig. 4A). The average size of clusters is controlled by varying a hyper-parameter called resolution (Fig. S8A). Systematically varying clustering resolution in our dataset revealed that not only did low-resolution clusters delineate proteins belonging to the same organelles (Fig. 4A-B), clustering at higher resolution also enabled to delineate functional pathways and even molecular complexes of interacting proteins (Fig. 4A-C). This demonstrates that the spatial distribution of each protein in the cell is highly specific, to the point that proteins sharing closely related functions can be identified on the sole basis of the similarity between their spatial distributions. This is further illustrated by how finely high-resolution clusters encapsulate proteins specialized in defined cellular functions (Fig. 4C). For example, our analysis not only separated P-body proteins (cluster #83) from other forms of punctated cytoplasmic structures, but also unambiguously differentiated vesicular trafficking pathways despite their very similar localization patterns: the endosomal machinery (#40), plasma membrane endocytic pits (#117) or COP-II vesicles (#143) were all delineated with high precision (Fig. 4C). Among ER proteins, the translocon clusters with the SRP receptor, EMC subunits and the OST glycosylation complex, all responsible for co-translational operations (#9). This performance extends to cytoplasmic (Fig. S8A) and nuclear clusters (Fig. S8B), revealing that spatial patterning is not limited to membrane-bound organelles and that sub-compartments exist also in the nucleo-cytoplasm. An illustrative example is a cytoplasmic cluster (#17) formed by a group of RNA-binding proteins (including ATXN2L, NUFIP2 or FXR1, Fig. 4C) that separate into granules upon stress conditions (*56*–*59*). Stress granules are not formed under the standard growth conditions used in our experiments, but the ability of our analysis to cluster these proteins together reveals an underlying specificity to their cytoplasmic localization (i.e., “texture”) even in the absence of stress.

**Figure 4:**
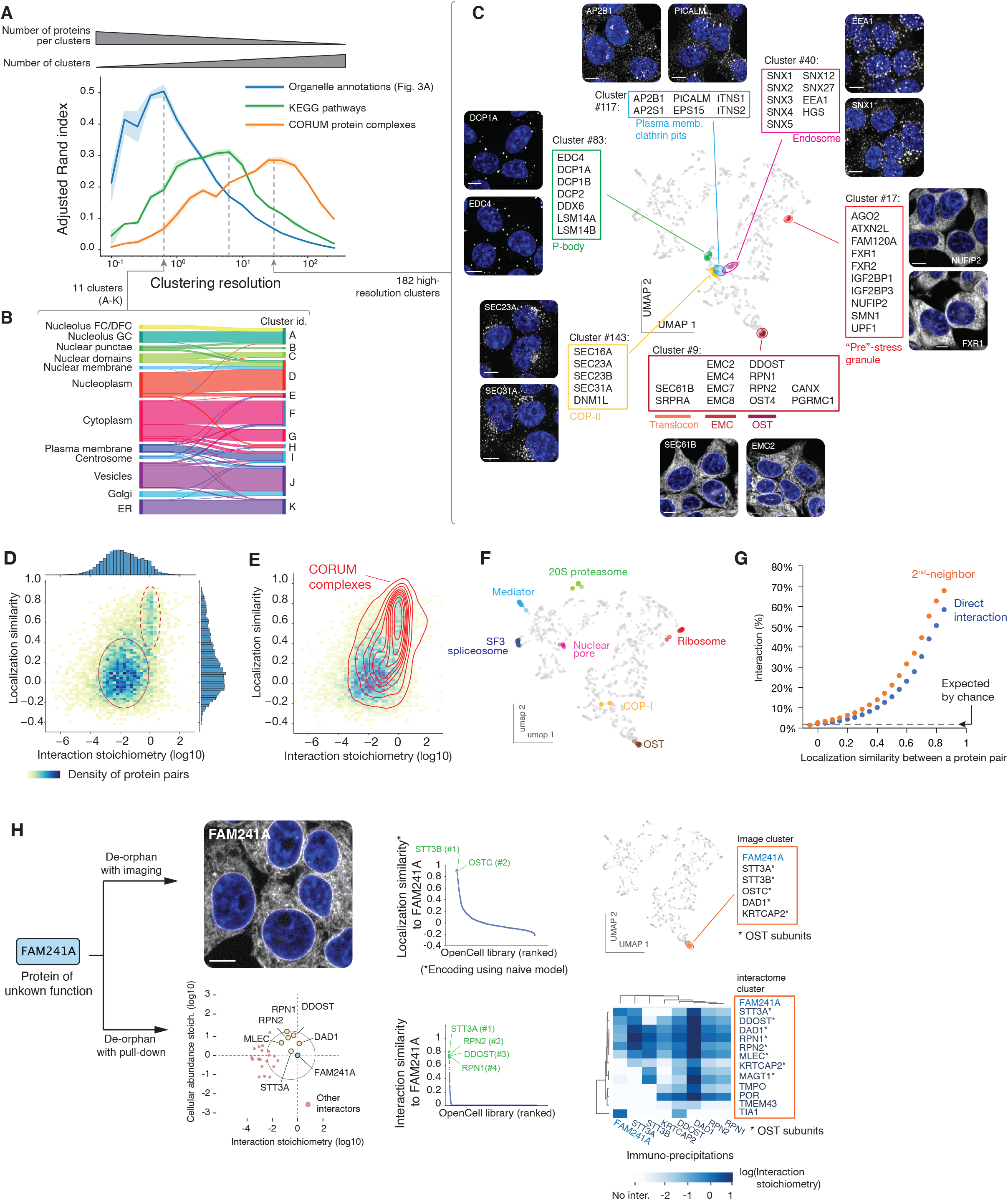
protein functional features derived from unsupervised image analysis. **(A)** Comparison of image-based Leiden clusters with ground-truth annotations. The Adjusted Rand Index (ARI, (86)) of clusters relative to three ground-truth datasets is plotted as a function of the Leiden clustering resolution. ARI (a metric between 0 and 1, see Materials and Methods) measures how well the groups from a given partition (in our case, the groups of proteins delineated at different clustering resolutions) match groups defined in a reference set. The amplitude of the ARI curves is approximately equal to the number of pairs of elements that partition similarly between sets; the resolution at which each curve reaches its maximum corresponds to the resolution that best captures the information in each ground-truth dataset. At a low resolution, Leiden clustering delineates groups that recapitulate about half of the organellar localization annotations, while at increasing resolutions, clustering recapitulates about a third of pathways annotated in KEGG, or molecular protein complexes annotated in CORUM. Shaded regions show standard deviations calculated from 9 separate repeat rounds of clustering, and average values are shown as a solid line. **(B)** High correspondence between low-resolution image clusters and cellular organelles. **(C)** Examples of functional groups delineated by high-resolution image clusters, highlighted on the localization UMAP. **(D)** Heatmap distribution of localization similarity (defined as the Pearson correlation between two deep learning-derived encoding vectors) vs. interaction stoichiometry between all interacting pairs of OpenCell targets. Two discrete sub-groups are outlined: low stoichiometry/low localization similarity pairs (solid line) and high stoichiometry/high localization similarity pairs (dashed line). **(E)** Probability density distribution of CORUM interactions mapped on the graph from (D). Contours correspond to iso-proportions of density thresholds for each 10th percentile. **(F)** Localization patterns of different subunits from example stable protein complexes, represented on the localization UMAP. **(G)** Frequency of direct (1st-neighbor) or once-removed (2nd neighbor, having a direct interactor in common) protein-protein interactions between any two pairs of OpenCell targets sharing localization similarities above a given threshold (x-axis). **(H)** Parallel identification of FAM241A as a new OST subunit by imaging or mass-spectrometry. See text for details.

A direct comparison between imaging and interactome data allows us to further examine the extent to which molecular-level relationships (that is, protein interactions) can be derived from a comparison of localization patterns. For OpenCell targets that directly interact, we compared the correlation between their localization encodings derived from machine learning (defining a “localization similarity”) and the stoichiometry of their interaction. This “localization similarity” measures the similarity between the global steady-state distributions of two proteins, as opposed to a direct measure of co-localization. We find that most proteins interact with low stoichiometry (as we previously described (*7*)) and without strong similarities in their spatial distribution (Fig 4D, solid oval). This means that while low-stoichiometry interactors co-localize at least partially to interact, their global distribution within the cell is different at steady state. On the other hand, high stoichiometry interactors share very similar localization signatures (Fig 4D, dashed oval). Indeed, proteins interacting within stable complexes annotated in CORUM fall into this category (Fig 4E), and the localization signatures of different subunits from large complexes are positioned very closely in UMAP embedding (Fig. 4F). In an important correlate, we found that a high similarity of spatial distribution is a strong predictor of molecular interaction. Across the entire set of target pairs (predicted to interact or not), proteins that share high localization similarities are also very likely to interact (Fig. 4G). For example, target pairs with a localization similarity greater than 0.85 have a 58% chance of being direct interactors, and a 68% chance of being second-neighbors (i.e., sharing a direct interactor in common). This suggests that protein-protein interactions could be identified from a quantitative comparison of spatial distribution alone. To test this, we focused on FAM241A (C4orf32), a protein of unknown function that was not part of our original library and asked whether we could predict its interactions using imaging data alone, compared to the classical de-or-haning approach that uses interaction proteomics. We thus generated a FAM241A endogenous fusion that was analyzed with live imaging and IP-MS separately. Encoding its localization pattern using a “naïve” machine learning model that was never trained with images of this new target revealed a very high localization similarity with two subunits of the ER oligo-saccharyl transferase OST (>0.85 similarity to STT3B and OSTC), and high-resolution Leiden clustering placed FAM241A in an image cluster containing only OST subunits (Fig 4H, top). This analysis suggested that FAM241A is a high-stoichiometry interactor of OST. IP-MS identified that FAM241A was indeed a stoichiometric subunit of the OST complex (Fig. 4H, bottom). While the specific function of FAM241A in protein glycosylation remains to be fully elucidated, this proof-of-concept example establishes that live-cell imaging can be used as a specific readout to predict molecular interactions.

Collectively, our analyses establish that the spatial distribution of a given protein contains highly specific information from which precise functional attributes can be extracted by modern machine learning algorithms. In addition, we show that while high-stoichiometry interactors share very similar localization patterns, most proteins interact with low stoichiometry and share different localization signatures. This reinforces the importance of low-stoichiometry interactions for defining the overall structure of the cellular network, not only providing the “glue” that holds the interactome network together (*7*) but also connecting different cellular compartments.

## RNA-binding proteins form a unique group in both interactome and spatial networks

To gain insight into global signatures that organize the proteome, we further examined the structures of our imaging and interactome datasets. First, we reduced the dimensionality of each dataset by grouping proteins into their respective spatial clusters (as defined by the high-resolution localization-based clusters in Figs. 4A, 4C) or interaction communities (as defined in Fig. 2B). We then separately clustered these spatial groups (Fig. S9A) and interaction communities (Fig. S9B) to formalize paired hierarchical descriptions of the human proteome organization. These hierarchies are highly structured and delineate clear groups of proteins (see comparison to hierarchies expected by chance, Fig. S9C). In both hierarchies, groups isolated at an intermediate hierarchical layer outline “modules” which are enriched for specific cellular functions or compartments (Fig. S9A-B; full ontology analysis in Suppl. Tables 5 & 9). At a higher layer, each dataset is partitioned into three “branches”, which represent core signatures that shape the proteome’s architecture from a molecular or spatial perspective (Fig. S9A-B). The structure of the localization-based hierarchy (Fig. S9A) recapitulates the human cell’s architecture across its three key compartments (nucleus, cytoplasm, membrane-bound organelles, Fig. S10A-B), which validates the relevance of our unsupervised hierarchical analysis. This motivated a deeper examination of the hierarchical architecture of the interactome (Fig. S9B, ontology analysis in Table S5). We found that intermediate-layer modules of the interactome delineate specific cellular functions such as transcription or vesicular transport (Fig. S9B), reflecting as expected that functional pathways are formed by groups of proteins that physically interact (*60*, *61*). More strikingly, the highest-layer structure showed that two of the three interactome branches were defined by clear functional signatures (Fig. S10C-E): branch B is significantly enriched in proteins that reside in or interact with lipid membranes, while branch C is significantly enriched in RNA-binding proteins (RNA-BPs) (Fig. 5B). This indicates that both membrane-related proteins and RNA-BPs interact more preferentially with each other than with other kinds of proteins in the cell.

**Figure 5:**
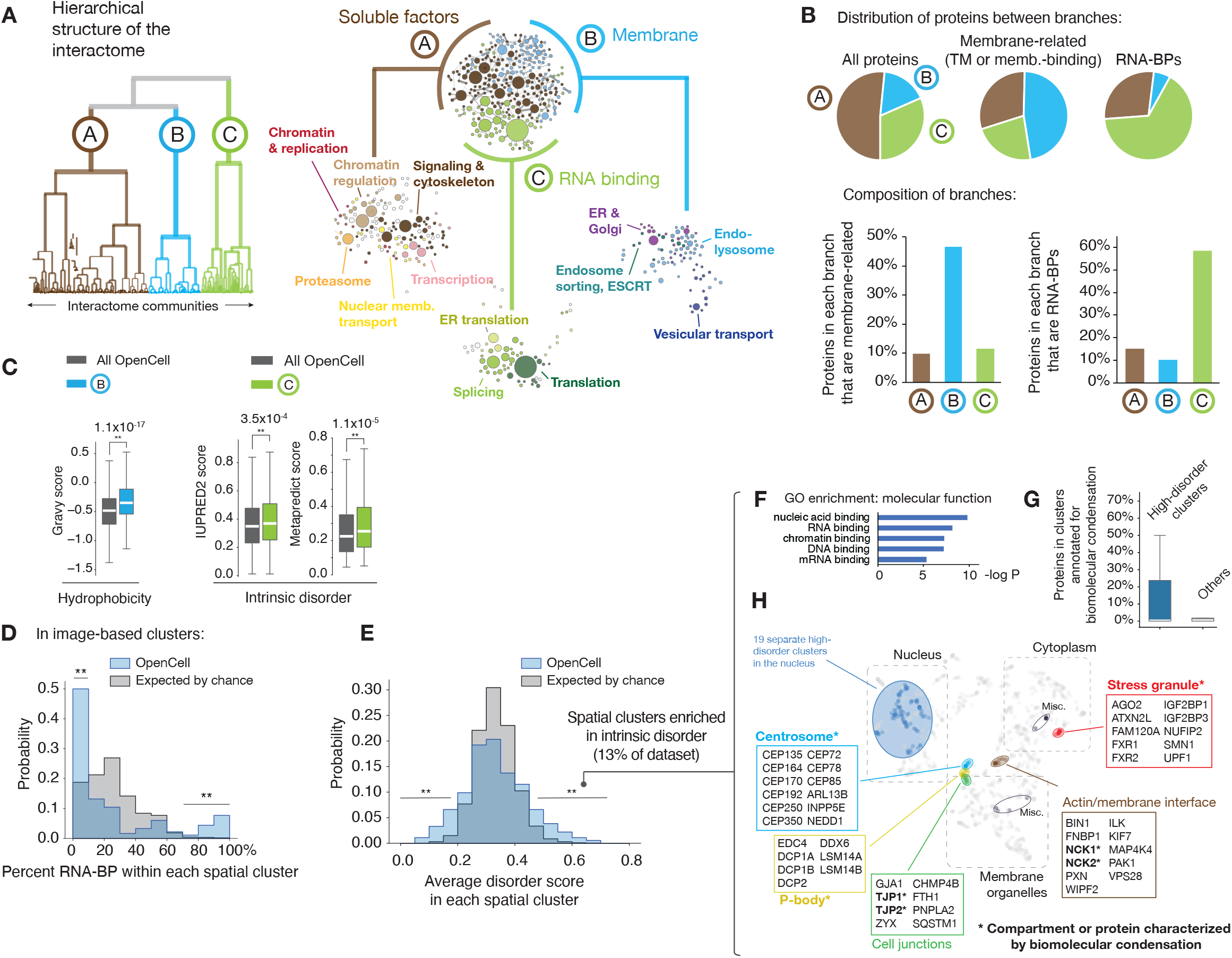
segregation of RNA-BPs in both interactome and imaging datasets. **(A)** Hierarchical structure of the interactome dataset, see full description in Figure S9B. **(B)** Distribution of membrane-related (transmembrane or membrane-binding) and RNA-BPs within the three interactome branches. **(C)** Distribution of intrinsic disorder in the RNA-BP branch of the interactome hierarchy (related to Figure S10). Two separate scores are shown for completeness: IUPRED2 (87), and metapredict (88), a new aggregative disorder scoring algorithm. Boxes represent 25th, 50th, and 75th percentiles, and whiskers represent 1.5x inter-quartile range. Median is represented by a white line. ** p < 10-4 (Student’s t-test), exact p-values are shown. **(D)** Distribution of RNA-BP percentage across spatial clusters, comparing our data to a control in which the membership of proteins across clusters was randomized 1,000 times. Lines indicate parts of the distribution over-represented in our data vs control (**: p < 2×10^−3^, Fisher’s exact t-test). **(E)** Distribution of disorder score (IUPRED2) across spatial clusters, comparing our data to a control in which the membership of proteins across clusters was randomized 1,000 times. Lines indicate parts of the distribution over-represented in our data vs control (**: p < 2×10^−3^, Fisher’s exact t-test). **(F)** Ontology enrichment analysis of proteins contained in high-disorder spatial clusters (average disorder score > 0.45). Enrichment compares to the whole set of OpenCell targets (p-value: Fisher’s exact test). **(G)** Prevalence of proteins annotated to be involved in biomolecular condensation in high-disorder vs. other spatial clusters. Boxes represent 25th, 50th, and 75th percentiles, and whiskers represent 1.5x inter-quartile range. Median is represented by a white line. Note that for both distributions, the median is zero. **(H)** Distribution of high-disorder spatial clusters in the UMAP embedding from Fig. 3D. Individual nuclear clusters are not outlined for readability. Multiple high-disorder spatial clusters include compartments or proteins known to be characterized by biomolecular condensation behaviors, which are marked by an asterisk.

That membrane-related proteins form a specific interaction group is perhaps not surprising as the membrane surfaces that sequester them within the three-dimensional cell will be partially maintained upon detergent solubilization. On the other hand, the fact that RNA-BPs also form a specific interaction group is unexpected, since our protein interactions were measured in nuclease-treated samples (*21*) in which most RNAs are degraded. This suggests that protein features beyond binding to RNAs themselves might drive the preferential interactions of RNA-BPs with each other. Therefore, we reasoned that the biophysical properties of proteins within each interactome branch might underly their segregation. Indeed, an analysis of protein sequence features revealed a separation of different biophysical properties in each branch (Fig. S10F-G). Branch B was enriched for hydrophobic sequences (Fig. 5C), consistent with its enrichment for membrane-related proteins, while branch C was enriched for intrinsic disorder (Fig. 5C). This is consistent with the fact that RNA-BPs are significantly more disordered than other proteins in the proteome (Fig. S11A, (*62*)). RNA-BPs are also among the most abundant in the cell (Fig. S11B), and form a higher number of interactions than other proteins (Fig. S11C-D).

IP-MS measures protein interactions *in vitro* after lysis and therefore does not directly address the spatial relationship between interacting proteins. Thus, we sought to further examine how RNA-BPs distribute in our live-cell imaging data. If RNA-BPs segregate into interacting groups *in vivo*, this should also manifest at the level of their intracellular localization: they should enrich in the same spatial clusters derived from our unsupervised machine learning analysis. Indeed, the distribution of RNA-BP content within spatial clusters revealed a significant over-representation of clusters that are either strongly enriched or depleted for RNA-BPs (Fig. 5D). Since spatial clusters can be interpreted as defining “micro-compartments” within the cell, both enrichment and depletion have functional implications: not only are RNA-BPs enriched within the same microcompartments, they tend to also be excluded from others. 16 out of the 26 spatial clusters (62%) that are highly enriched in RNA-BPs include at least one protein involved in biomolecular condensation (as curated in PhaSepDB (*63*)), which might reflect a prevalent role for biomolecular condensation in shaping the RNA-BP proteome. Collectively, both interactome and imaging data underscore that RNA-BPs (a prevalent group of proteins that represents 13% of proteins expressed in HEK293T cells, see Table S2) form a distinct sub-group within the proteome characterized by unique properties.

These results motivated a broader analysis of the contribution of intrinsic disorder to the spatial organization of the proteome in our dataset. Plotting the distribution of mean intrinsic disorder within spatial clusters revealed a significant over-representation of clusters both enriched and depleted in disordered proteins (Fig. 5E). 26 out of 182 total spatial clusters were enriched for disordered proteins, covering 13% of the proteins in our imaging dataset. Overall, the extent to which disordered proteins segregate spatially is similar to the degree of segregation found for hydrophobic proteins: an analogous analysis revealed that 10% of proteins in our dataset are found within clusters significantly enriched for high hydrophobicity (Fig. S12E), which map to membrane-bound organelles (Fig. S12F). This supports the hypothesis that intrinsic disorder is as important a feature as hydrophobicity in organizing the spatial distribution of the human proteome. Consistent with our previous analysis, high-disorder clusters were enriched for RNA-BPs (Fig. 5F), with 15 out of these 26 clusters containing over 50% of RNA-BPs. High-disorder clusters were also enriched for proteins annotated to participate in biomolecular condensation (Fig. 5G), and were predominantly found in the nucleus (19 clusters, 73% of total, Fig. 5H). 5 out of 7 high-disorders clusters found in the cytosol delineate compartments for which biomolecular condensation has been proposed to play an important role (Fig. 5G), namely P-bodies (*64*), stress granules (*59*), centrosome (*65*), cell junctions (*66*) and the interface between cell surface and actin cytoskeleton (*67*).

## Interactive data sharing at opencell.czbiohub.org

To enable widespread access to the OpenCell datasets, we built an interactive web application that provides side-by-side visualizations of the 3D confocal images and of the interaction network for each tagged protein, together with RNA and protein abundances for the whole proteome (Fig. 6). Our web interface is fully described in Suppl. Fig S12.

**Figure 6:**
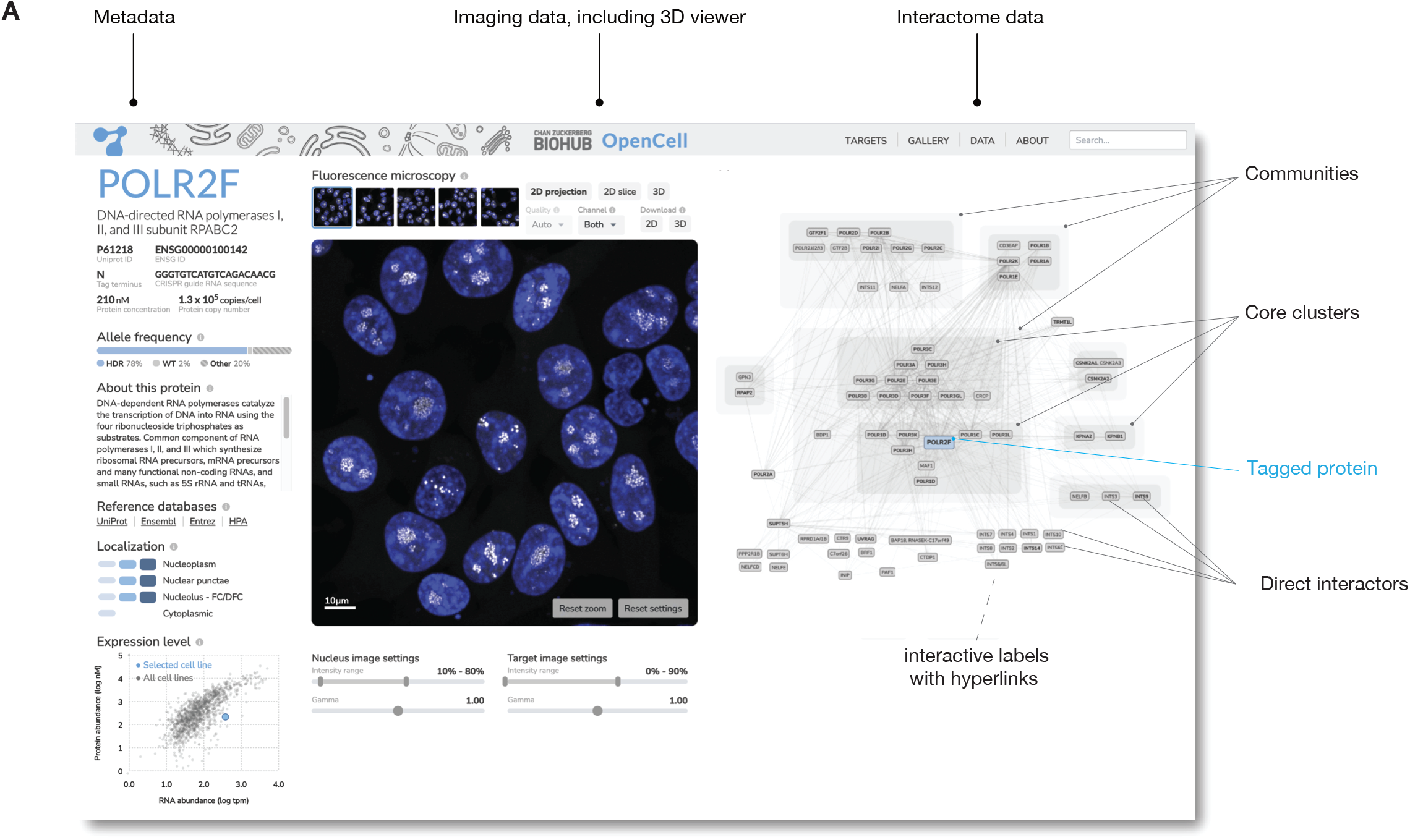
the OpenCell website. Shown is an annotated screenshot from our web-app at http://opencell.czbiohub.org, which is described in more details in Suppl Fig. S12.

## Discussion

OpenCell combines three strategies to augment the description of human cellular architecture. First, we present an integrated experimental pipeline for high-throughput cell biology, fueled by scalable methods for genome engineering, live-cell microscopy and IP-MS. Second, we provide an open-source resource of well-curated localization and interactome measurements, easily accessible through an interactive web interface at opencell.czbiohub.org. And third, we developed an analytical framework for the representation and comparison of interaction or localization signatures (including a self-supervised machine learning approach for image encoding). Finally, we demonstrate how our dataset can be used both for fine-grained mechanistic exploration (to explore the function of multiple proteins that were previously uncharacterized), as well as for investigating the core organizational principles of the proteome.

Our current strategy that combines split-FPs and HEK293T – a cell line that is heavily transformed but easily manipulatable – is mostly constrained by scalability considerations. Excitingly, technological advances are quickly broadening the set of cellular systems that can be engineered and profiled at scale. Advances in stem cell technologies enable the generation of libraries that can be differentiated in multiple cell types (*11*), while innovations in genome engineering (for example, by modulating DNA repair (*68*)) pave the way for the scalable insertion of gene-sized payload, for the combination of multiple edits in the same cell, or for increased homozygosity in polyclonal pools. In addition, recent developments in high-throughput light-sheet microscopy (*69*) might soon enable the systematic description of 4D intracellular dynamics (*70*).

A central feature of our approach is to use endogenous fluorescent tags to study protein function. Genome-edited cells enable to examine protein function at near-native expression levels (which can circumvent some limitations of over-expression (*71*)), and to measure protein localization in live cells (which can avoid artefacts caused by fixation or antibody labeling (*72*)). Comparing our data to the current reference datasets of protein-proteins interactions (Fig. S4C-F) or localization (Fig. S7C-D) highlights the performance of our strategy. In addition, our high success rate tagging essential genes (Fig. S2A; see also (*73*) in yeast) and the successful tagging of the near-complete yeast proteome (*14*, *73*) support that fluorescent tagging generally preserves normal protein physiology. However, limitations exist for specific protein targets. FPs are as big as an average human protein and their insertion can impair function or localization, for example by occluding important interaction interfaces or impairing sub-cellular targeting sequences. In other cases, tags can affect expression or degradation rates, which might explain why we find tagged proteins being expressed at 80% of their endogenous abundance, and 8% of targets in our dataset having outlier abundances at steady-state (Fig. S3D). Further, tagging often cannot discriminate between different isoforms of a protein (such as splicing or post-translationally modified variants). Finally, relying on endogenous expression can be an obstacle given the low concentration of most proteins in the human cell: even using a very bright FP like mNeonGreen (*74*), detecting proteins in the bottom 50% percentile of abundance is difficult (Fig. S2D). Solutions to this obstacle include using FP repeats to increase signal (*18*, *23*) or using tags that bind chemical fluorophores (e.g., HaloTag (*75*)), which can be brighter than FPs or operate at wavelengths where cellular auto-fluorescence is decreased (*76*). Overall, the full description of human cellular architecture remains a formidable challenge which will require complementary methods being applied in parallel. The diversity of large-scale cell biology approaches is a solution to this problem (*6*, *8*, *9*, *11*, *31*, *70*, *77*–*80*). Mirroring the advances in genomics following the human genome sequence (*2*), opensource systematic datasets will likely play an important role in how the growth of cell biology measurements can be transformed into fundamental discoveries by an entire community (*81*).

In addition to presenting a resource of measurements and protocols, we also demonstrate how our data can be used to study the global signatures that pattern the proteome. Our analysis reveals that RNA-binding proteins, which form one of the biggest functional family in the cell, are characterized by a unique set of properties and segregate from other proteins in term of both interactions and spatial distribution. It would be fascinating to explore to which extent RNA itself might act as a structural organizer of the cellular proteome (*62*, *82*). This is for example the case for some non-coding RNAs whose main function is to template protein interactions to form nuclear bodies (*83*). High intrinsic disorder is one of the distinguishing features of RNA-BPs, which likely contributes to their unique properties. Beyond RNA-BPs, our data supports a general role for intrinsic disorder in shaping the spatial distribution of human proteins. For example, 13% of proteins in our dataset are found in spatial clusters that are significantly enriched for disordered proteins. This adds to the growing appreciation that intrinsic disorder, which is much more prevalent in eukaryotic vs. prokaryotic proteomes (*84*, *85*), plays a key role in the functional subcompartmentalization of the eukaryotic nucleo- and cytoplasm in the context of biomolecular condensation (*86*).

Lastly, we show that the spatial distribution of each human protein is very specific, to the point that remarkably detailed functional relationships can be inferred on the sole basis of similarities between localization patterns – including the prediction of molecular interactions (which complements other studies (*87*)). This highlights that intracellular organization is defined by fine-grained features that go beyond membership to a given organelle. Our demonstration that self-supervised deep learning models can identify complex but deterministic signatures from light microscopy images opens exciting avenues for the use of imaging as an information-rich method for deep phenotyping and functional genomics (*51*). Because light microscopy is easily scalable, can be performed live and enables measurements at the single-cell level, this should offer rich opportunities for the full quantitative description of cellular diversity in normal physiology and disease.

## Acknowledgements

We thank N. Neff, M. Tan, R. Sit and A. Detweiler for help with high-throughput sequencing; G. Margulis, A. Sellas, E. Ho and J. Mann for operational support; A. McGeever for help with web application architecture and deployment; and S. Schmid for critical feedback. M.D.L. thanks C. L. Tan for continuous discussions. H.K. was supported by an International Research Fellowship from the Japan Society for the Promotion of Science. R.M.B. was supported by a NIH pre-doctoral fellowship (F31 HL143882). B.H. was supported by NIH (R01GM131641) and is a Chan Zuckerberg Biohub Investigator. J.S.W. was supported by NIH (1RM1HG009490) and is an investigator with the Howard Hughes Medical Institute.

## Material and Methods

are available as a supplementary file.

---

**Figure S1:**
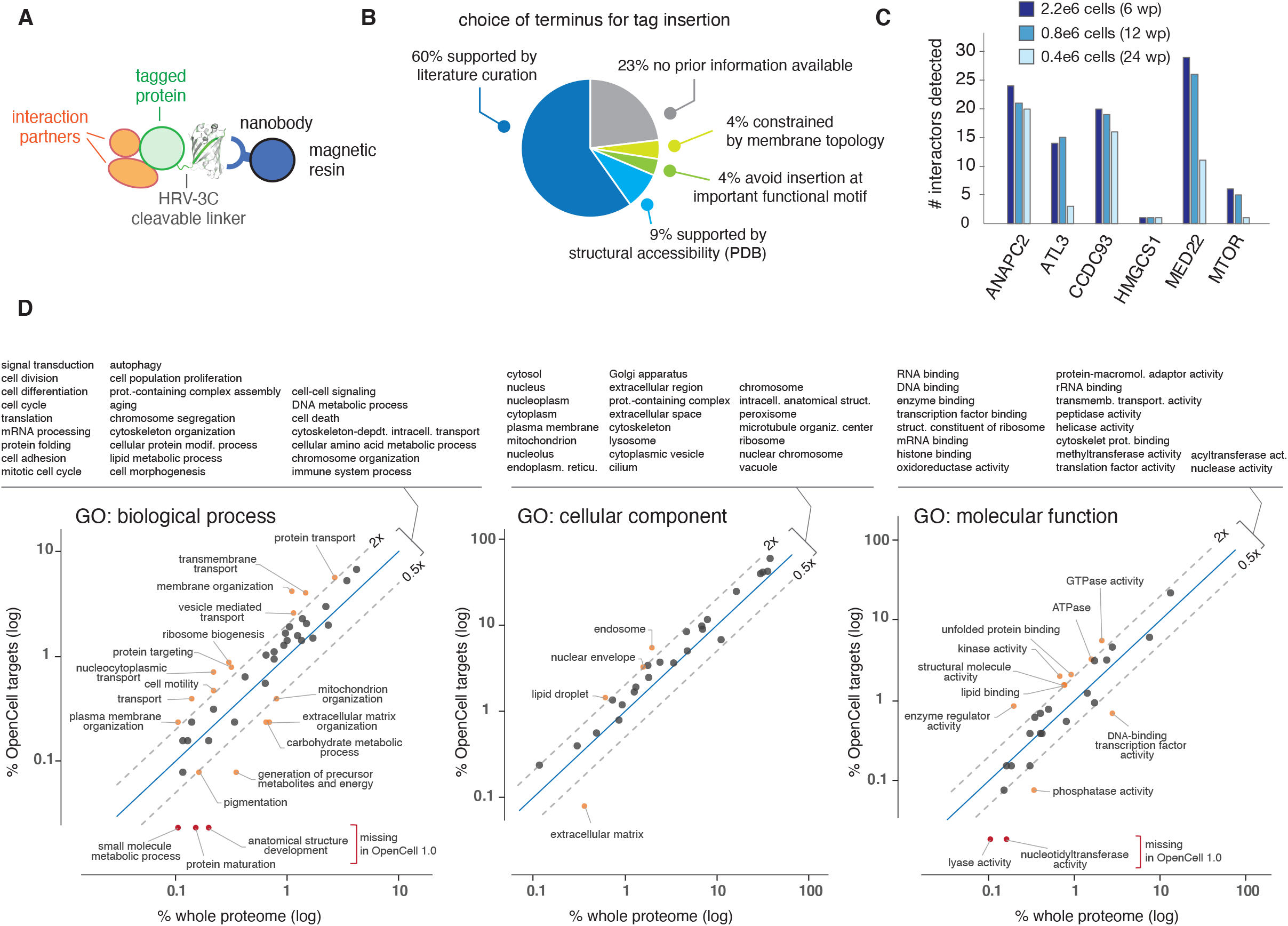
experimental pipeline (related to Figure 1). **(A)** IP/MS using FP capture. All mNG11 tagging constructs also include an HRV-3C cleavable linker for optional release from the capture resin. **(B)** Justifying the choice of tag insertion in engineered cell lines. To inform tag insertion sites, we used a combination of existing data from the literature suggesting preservation of properties, 3D structures of protein complexes from the PDB and sequence analysis to avoid important functional motifs. 4% of insertion sites were constrained by the topology of transmembrane protein targets (fusion to cytosolic termini), and for 23% of targets no prior data was available. See details in Suppl. Table 1. **(C)** Sensitivity of interaction proteomics detection on a timsTOF instrument. The number of interactors detected in pull-downs from 6 different targets is shown, varying the amount of input material. To balance sensitivity and scalability, 0.8e6 cells were used for high-throughput assays (12 well-plate, wp). **(D)** Distribution of gene ontology annotations in the OpenCell library (successful targets only) compared the whole proteome. Over- and under-represented terms are outlined. Because organellar organization and transport between organelles are foundational to human cellular architecture, proteins in these groups are slightly enriched in our library. Under-represented groups are mostly comprised of proteins in compartments that are not accessible to our tagging strategy (mitochondrial functions, extracellular matrix) or proteins that are typically present at low copy numbers and therefore difficult to detect at endogenous levels (transcription factors).

**Figure S2:**
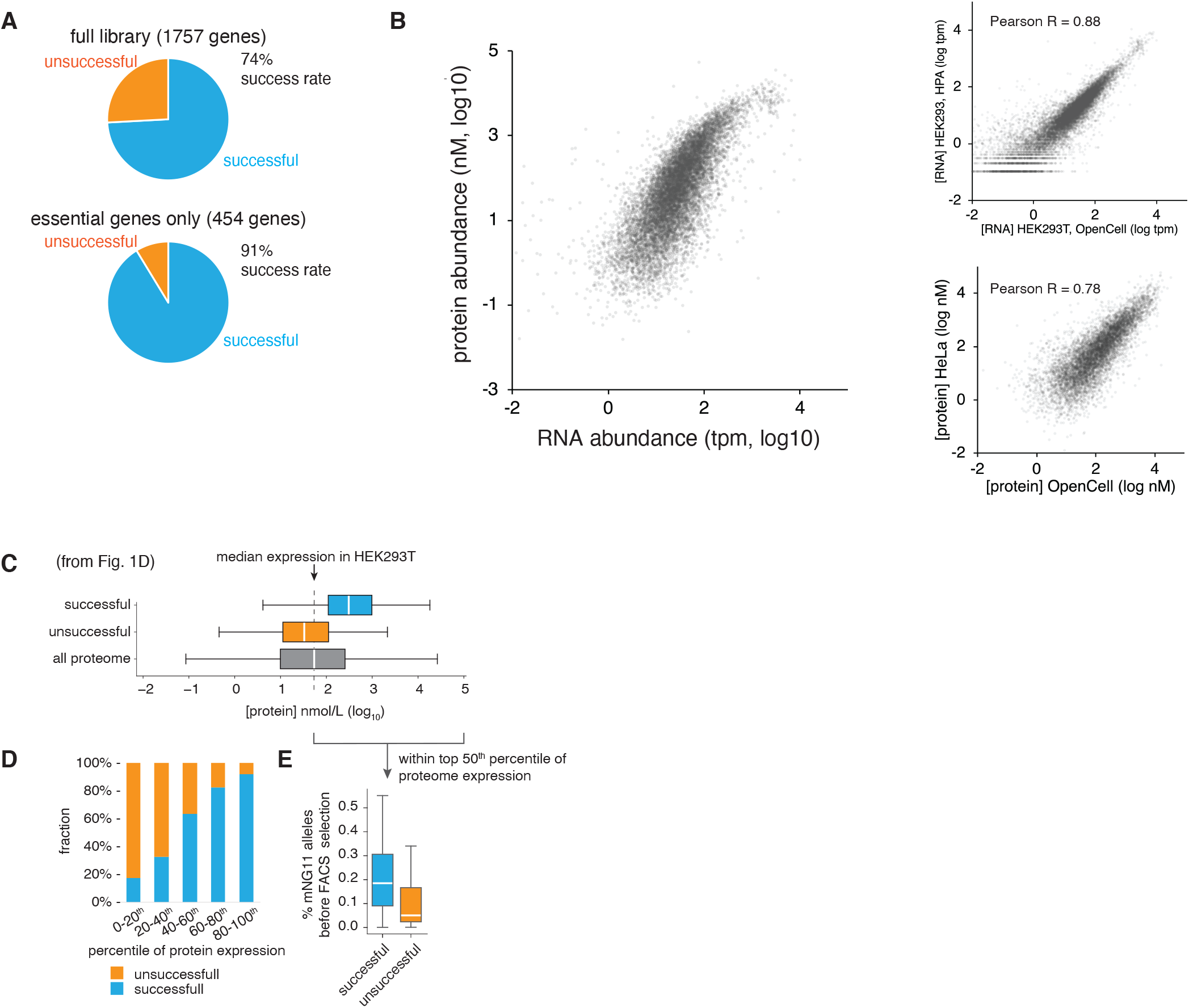
cell line generation (related to Figure 1). **(A)** Success rate for the generation and detection of fluorescently tagged cell lines are compared for the whole set of targets we attempted, and the subset of these that are essential genes. **(B)** Correlation of protein and RNA abundance in HEK293T cells (OpenCell). For comparison purposes, RNA and protein abundances in our dataset are compared to two external references: HEK293 cell line RNASeq from the Human Protein Atlas, and the HeLa proteome published in (7). In both cases, our data correlates well with existing references. **(C)** Repeated from Figure 1C. **(D)** Fluorescent detection success rates for proteins at different percentiles of abundance in the proteome. **(E)** For well-expressed proteins (top 50th percentile of abundance), successful detection is correlated with high rates of CRSIPR-mediated homologous recombination.

**Figure S3:**
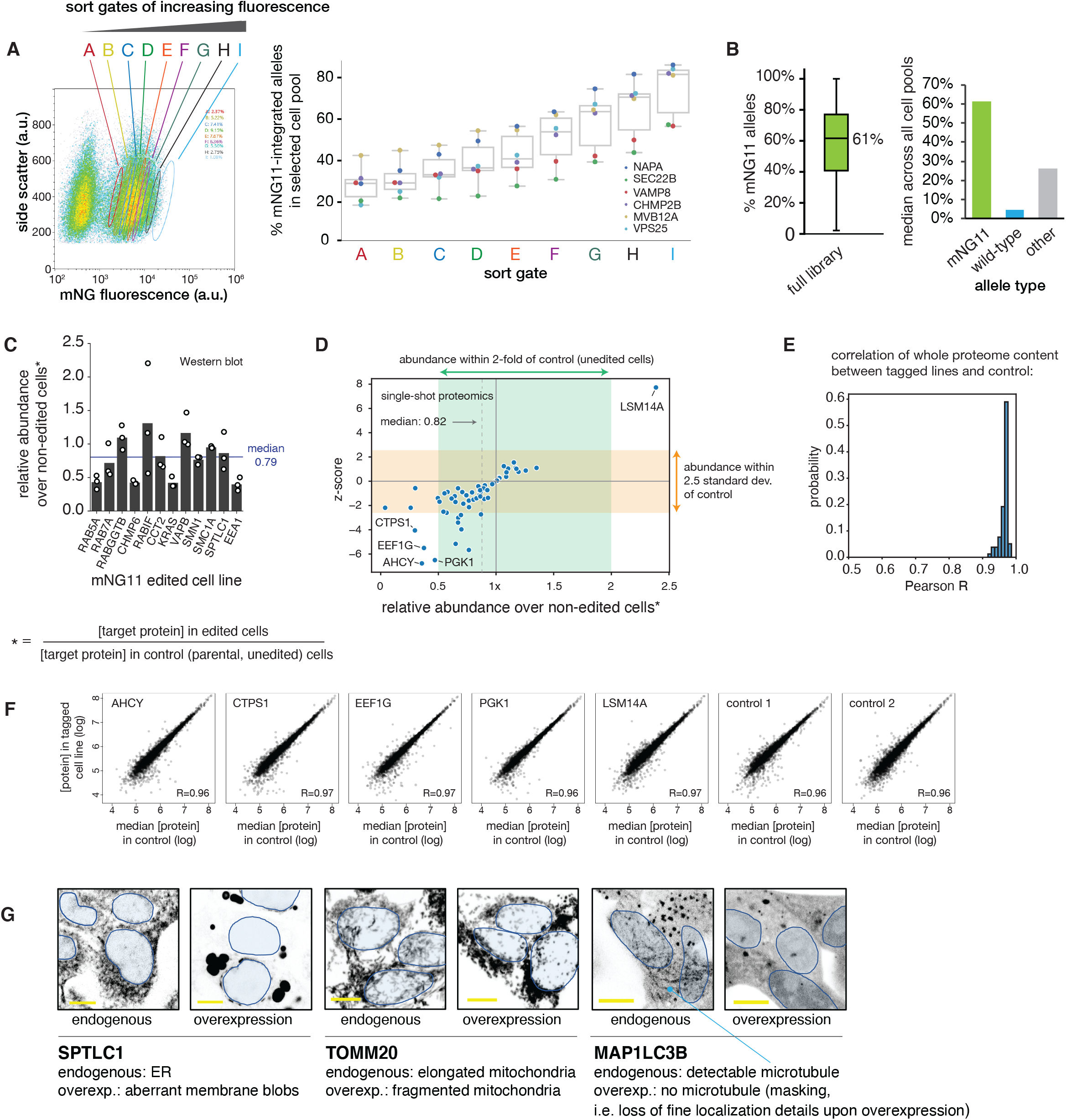
cell library characterization and quality control (related to Figure 1). **(A)** Optimization of sorting strategy. Polyclonal cell pools were sorted using gates of increasing fluorescence (left panel) and genotyped to quantify the enrichment for mNG11-inserted alleles (right panel, showing data for 6 different target genes). This informed our final sorting strategy in which the top 1% of fluorescent cells (gate I) were selected. **(B)** Genotype analysis of the polyclonal OpenCell library. A single allele is required for fluorescence, but our cell collection is enriched for homozygous insertions. In total, mNG11 insertions account for 61% (median) of alleles in a given cell pool across the full library (Boxes represent 25th, 50th, and 75th percentiles, and whiskers represent 1.5x interquartile range). The median values of mNG11 integrated alleles, wt alleles and other alleles are shown on the right. **(C)** Measurement of target protein abundance in final selected cell pools vs. parental cell line, by quantitative Western blotting. **(D)** Measurement of target protein abundance in final selected cell pools vs. parental cell line, by single-shot mass spectrometry. In these experiments, tagged lines are measured in a single replicate and compared to 6 replicates of non-edited control cell lines. Outliers targets are defined by an abundance that deviates by more than 2.5 standard deviations and by more than 2-fold of their abundance in the controls. The 5 outlier lines are outlined. **(E)** Distribution of Pearson correlation values measuring the overall correlation of abundances for all cellular proteins in each tagged cell line vs. median control. **(F)** For the outliers outlined in (D), correlation of abundances for all cellular proteins in the tagged cell line vs. median control. The abundance correlations for two individual control repeats are shown for reference. **(G)** Examples of overexpression artifacts. Single z-slice confocal images are shown (scale bar: 10 μm). Endogenously tagged lines and their equivalent overexpression constructs were not imaged using the same laser power, so that signal intensities are not directly comparable. Nuclei are shown as blue outlines (nuclei can be located in a different z-plane than the one shown). “Masking effects” are defined as the loss of fine localization details upon overexpression.

**Figure S4:**
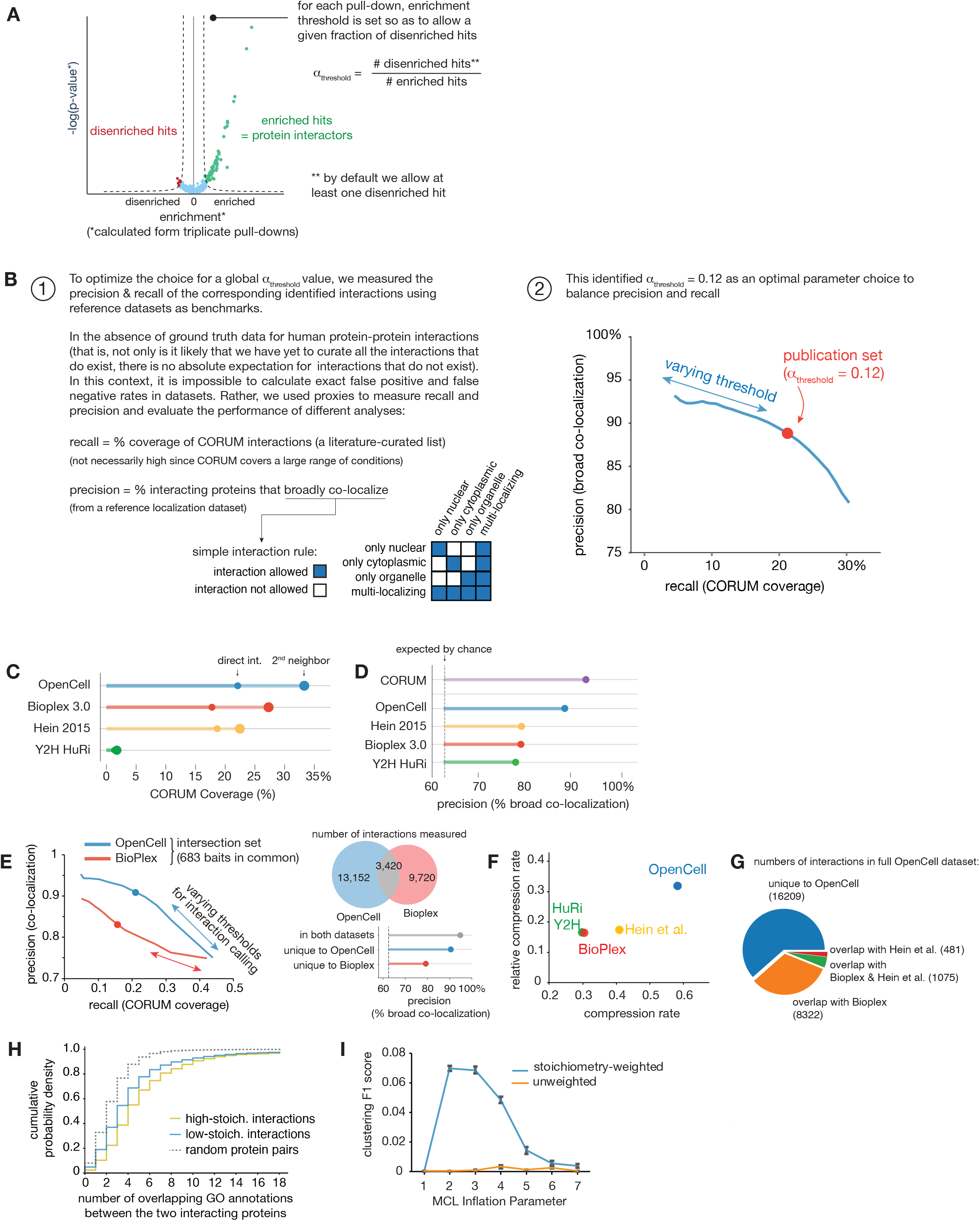
interactome analysis (related to Figure 2). **(A)** Strategy for defining enrichment threshold to define interactions. Our strategy builds upon methods described by Hein et al (7). Here we use a quantitative approach to define enrichment thresholds dynamically for each replicate set, globally constrained by the parameter α_threshold_. **(B)** To optimize parameter choice, we measured how precision (% co-localization) and recall (% CORUM coverage) of the corresponding interaction network varied with α_threshold_. This informed a final value of 0.12. **(C)** Comparing interaction recall (% CORUM coverage) of OpenCell vs. other large-scale interactomes, including direct or 2nd-neighbor interactions (i.e., sharing a direct interactor in common). **(D)** Comparing interaction precision (% co-localization) of OpenCell vs. other large-scale interactomes. CORUM interactions are shown as a reference. **(E)** Direct comparison of OpenCell vs. Bioplex 3.0 on identical bait set. Both datasets use the same HEK-293T cell line and share a large number (683) of baits in common. Precision and recall analysis by varying threshold for interaction detection (α_threshold_ in OpenCell and *pInt* in Bioplex) is shown for the intersection set of 683 baits (dots represent values using thresholds used for final publication sets in both studies). For these set of overlapping baits, OpenCell also includes many new measured interactions for that intersection set of baits (right panel, top). Interestingly, the interactions unique to Open-Cell have high precision values (right panel, bottom). **(F)** Compressibility analysis (31) of OpenCell vs. other large-scale interactomes. **(G)** Number of interactions measured in OpenCell (in the full dataset) that were also measured in Hein et al. (7) or BioPlex 3.0. **(H)** Distribution of GO annotation overlap between protein pairs identified in low-stoichiometry and high-stoichiometry interactions. **(I)** MCL clustering performance (F1 score) using stoichiometry-weighted or unweighted interaction graphs, derived from CORUM interactions as described in Drew et al (89).

**Figure S5:**
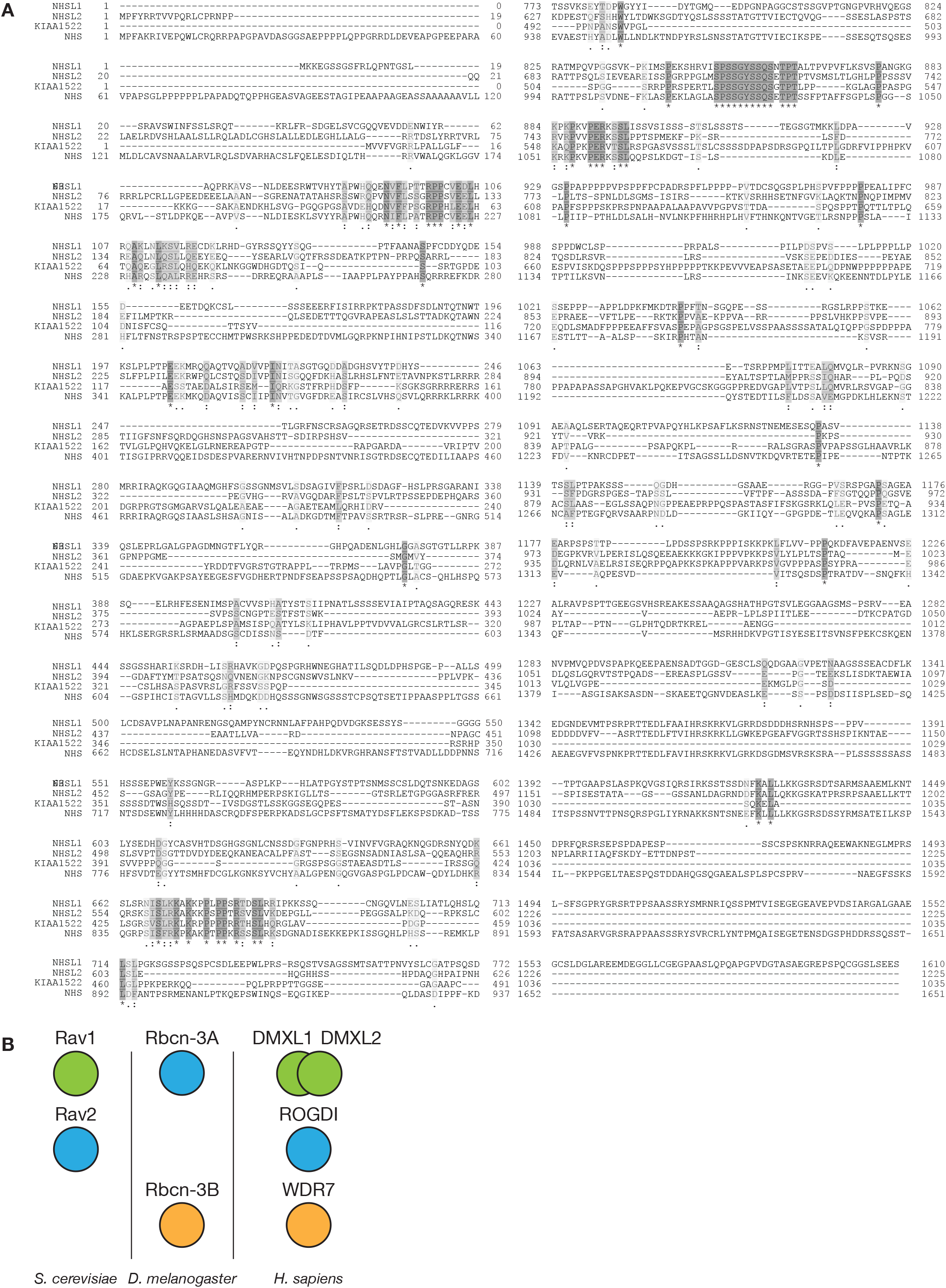
sequence analysis of orphan proteins (related to Figure 2). **(A)** Amino-acid sequence alignment between human NHSL1, NSHL2, KIAA1522 and NHS. **(B)** Correspondence of RAVE complex members in S. cerevisiae, D. melanogaster and H. sapiens. Note that in S. cerevisiae RAVE also includes Skp1, not depicted here.

**Figure S6:**
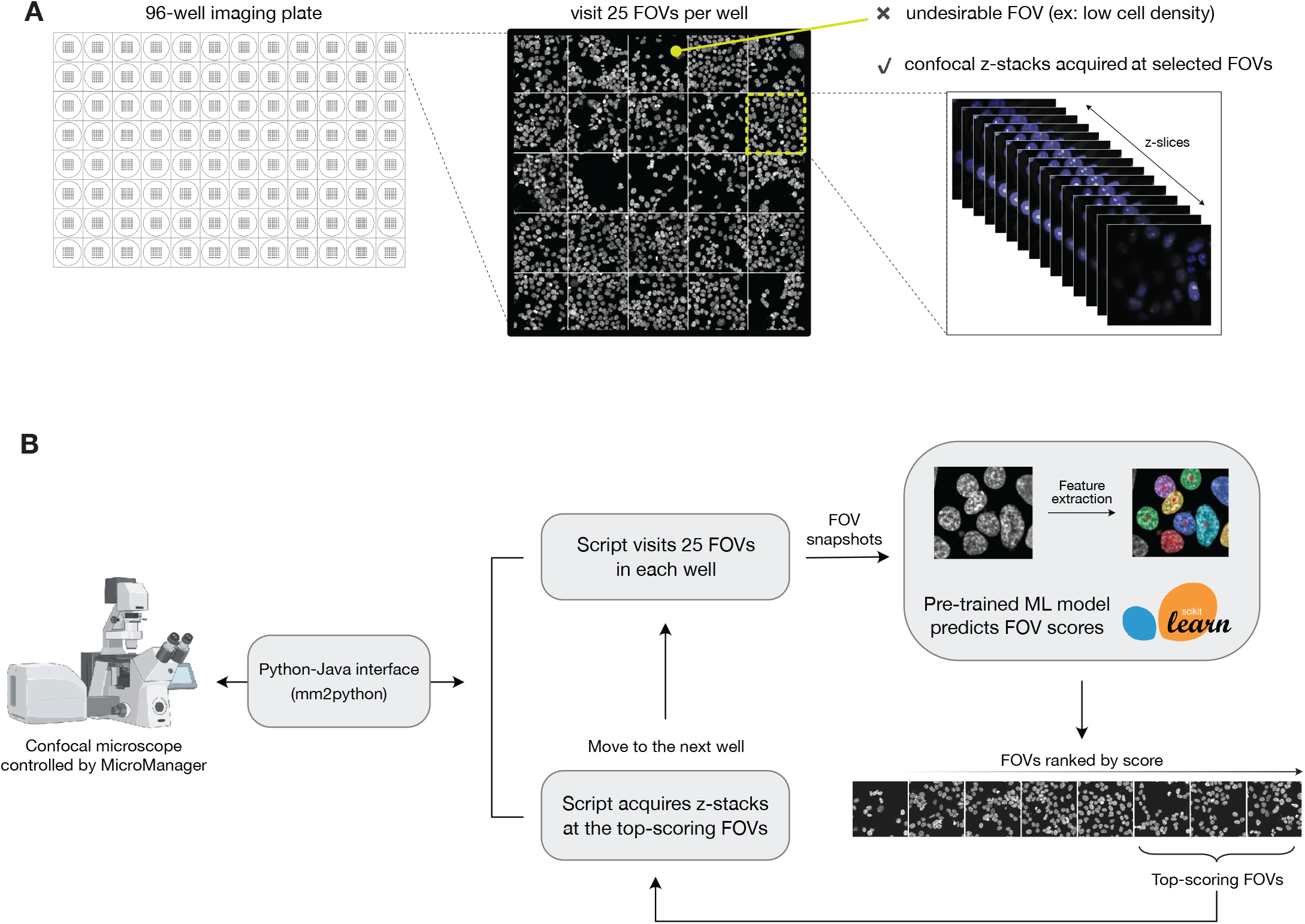
computer vision for automated microscopy acquisition (related to Figure 3). **(A)** To automate microscopy acquisition on 96-well plates and to limit experimental variability between imaging sessions (e.g., to limit variations in cell density) we paired an acquisition script, written in Python, with a pre-trained machine learning model to select field of views (FOVs) on-the-fly during the acquisition. A total of 25 FOVs are sampled per well in a single z-plane, and desirable FOVs are selected for further 3D confocal acquisition on the basis of a score predicted by the pre-trained model. **(B)** Microscopy automation workflow. Microscope hardware is controlled by a Python-based acquisition script via an open-source MicroManager-Python bridge (mm2python; https://github.com/czbiohub/mm2python). This approach enables us to combine custom acquisition logic with the rich ecosystem of Python-based machine-learning packages. Here, we use the scikit-image package to extract features from each FOV snapshot, then use a pre-trained random-forest regression model (scikit-learn) to predict a quality score for the FOV. This process is not computationally expensive and requires less than a second; the FOV score can therefore be used immediately to determine whether the script should acquire a z-stack or else move on to the next position. To maximize the quality of our confocal z-stacks, however, we chose to visit and score all 25 FOVs in each well, then re-visit the top-scoring FOVs for confocal z-stack acquisition.

**Figure S7:**
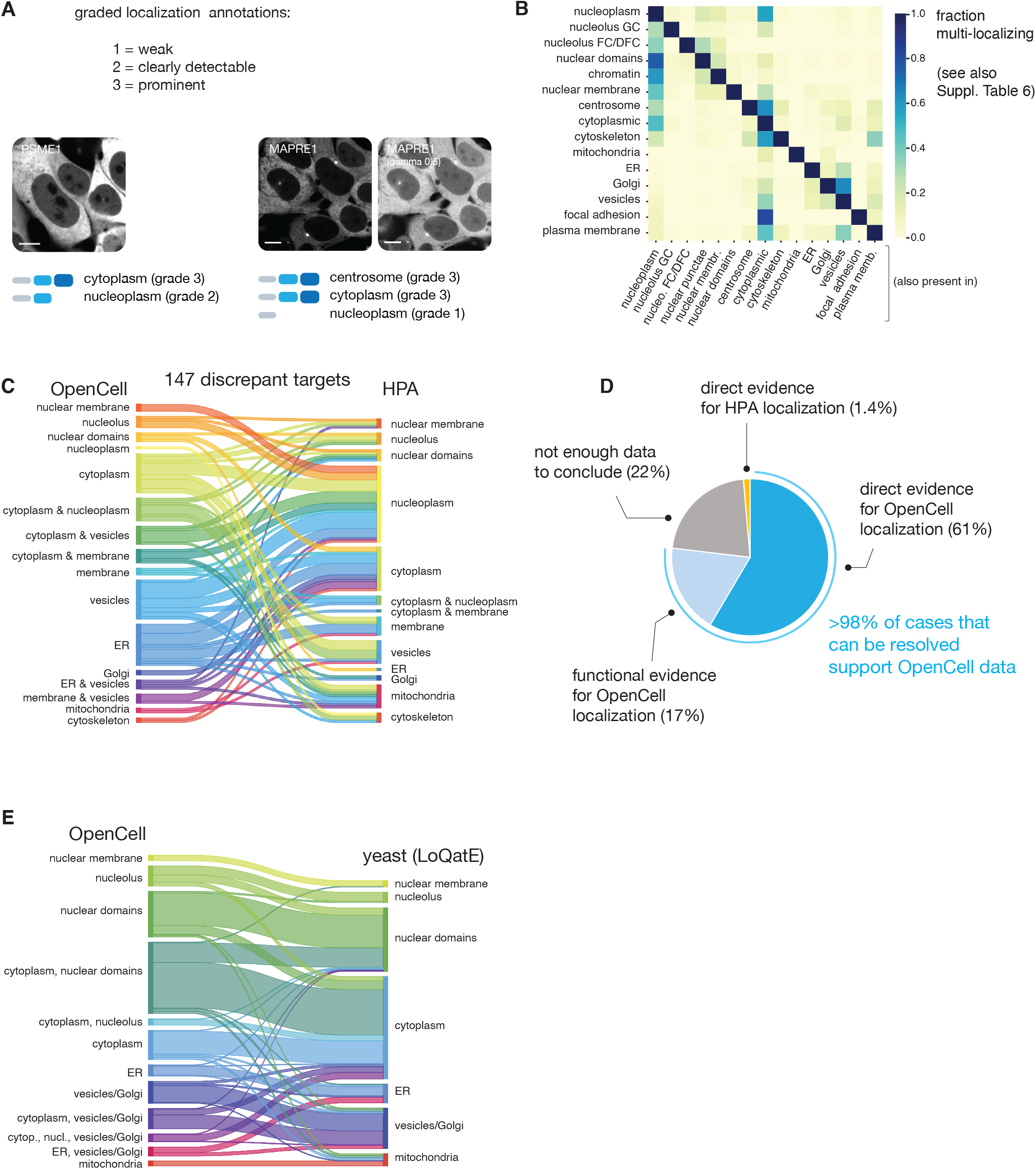
the OpenCell image dataset (related to Figure 3). **(A)** Principle of graded localization annotation (manual annotations). **(B)** Fraction of multi-localization between cellular compartments. Complete localization annotations can be found in Suppl. Table 6. **(C)** Comparison of annotated localization for proteins in OpenCell and Human Protein Atlas (HPA, version v20) datasets for which annotations are inconsistent. **(D)** Extensive literature curation allows to resolve 77% of OpenCell/HPA discrepancies (full details in Suppl. Table 8). Here “direct evidence” refers to proteins for which localization has been directly measured in published studies, while “functional evidence” refers to proteins for which localization might not have been directly measured, but for which literature establishes a function that is predictive of a specific localization. For example, SCFD1 is a protein whose main known function is to regulate transport between ER and Golgi. This qualifies as “functional evidence”. It is annotated as localized in the ER and Golgi in OpenCell, and in the nucleoplasm (main) and cytosol (additional) in HPA. **(E)** Comparison of annotated localization for 350 orthologous proteins in OpenCell and *S. cerevisiae* yeast (from LoQate (46)). Note that in yeast Golgi and vesicles are difficult to distinguish.

**Figure S8:**
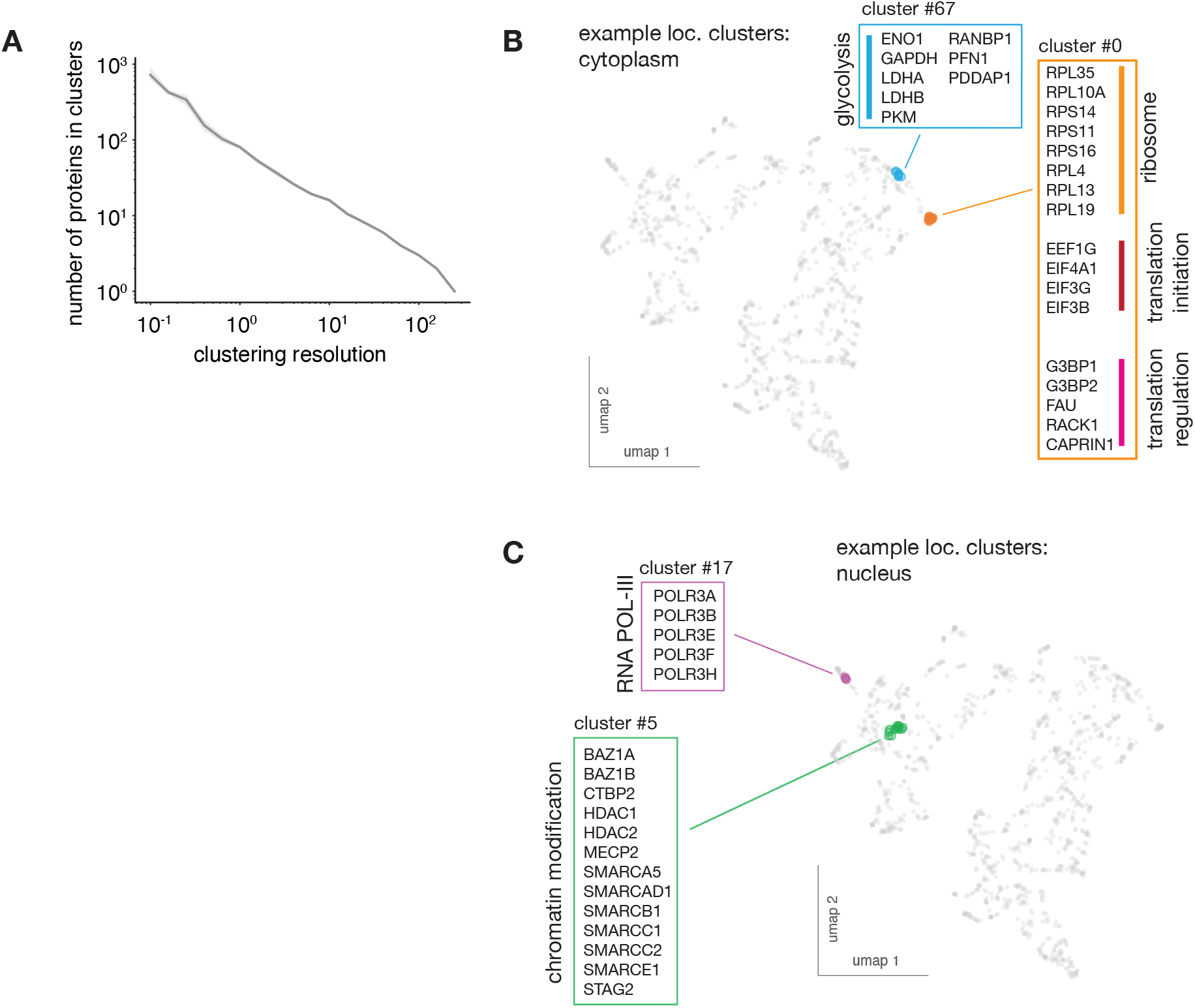
high-resolution image clusters (related to Figure 4C). **(A)** Size of clusters (number of proteins in each cluster) as a function of clustering resolution. Shaded regions show standard deviations calculated from 9 separate repeat rounds of clustering, and average values are shown as a solid line. **(B), (C)** Examples of clusters of cytoplasmic **(B)** and nuclear **(C)** proteins.

**Figure S9:**
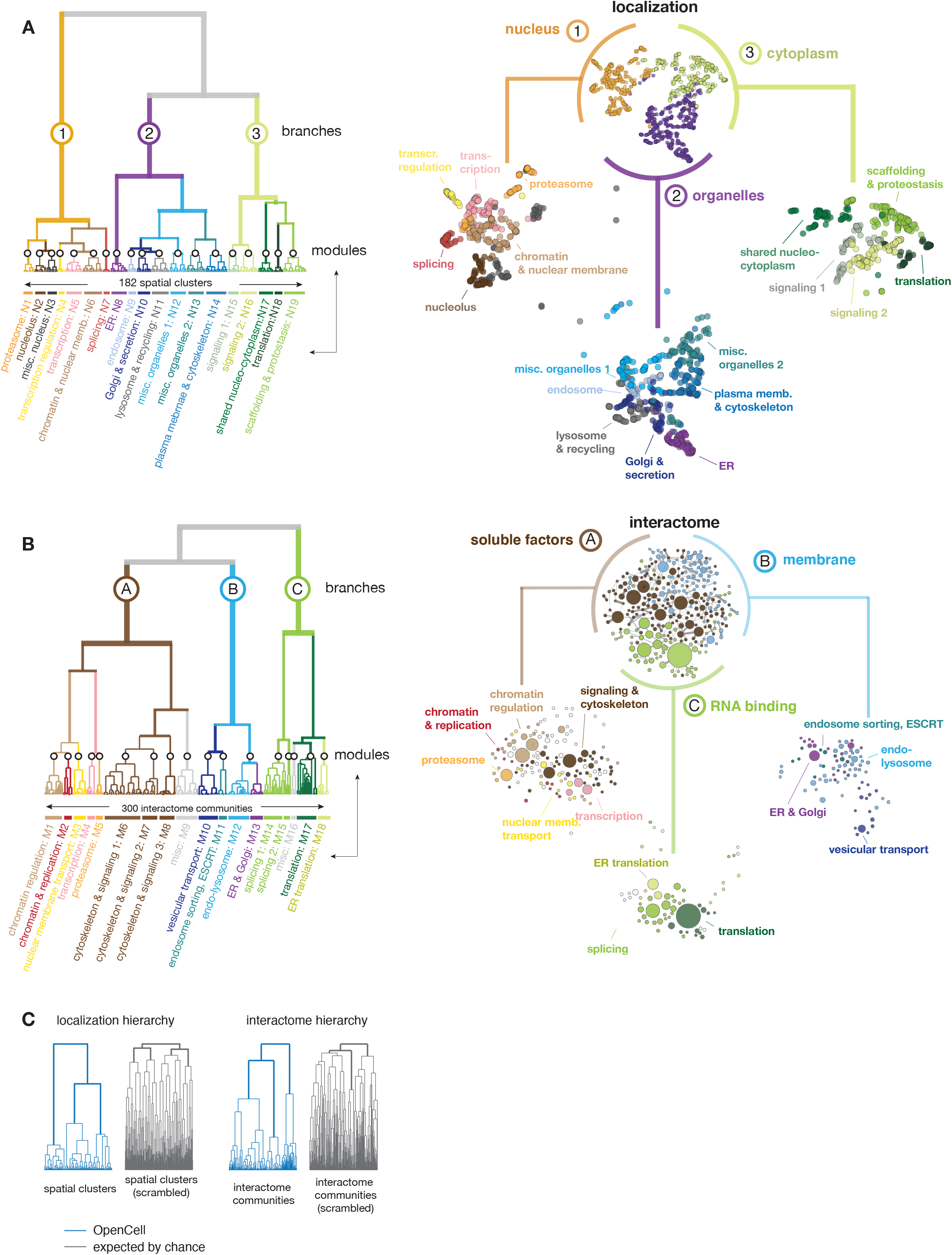
full hierarchical structure of interactome and localization datasets (related to Figure 5). Dendrograms represent the hierarchical relationships connecting **(A)** the full set of protein communities identified in the interactome (see Fig. 2) or **(B)** the full set of high-resolution clusters identified in the image collection (see Fig. 4C). For each dataset, an intermediate layer of hierarchy separates 18-19 modules, while an upper hierarchical layer delineates three separate branches. Modules and branches are annotated on the basis of gene ontology enrichment analysis (see Suppl. Tables 5 & 9). Right-hand panels present the topological arrangement of branches (top) and modules (bottoms) in each dataset, highlighted from the full graph of connections between interaction communities (“interactome”, see Fig. 2D) or from the localization UMAP (“localization”, see Fig. 4C). The color codes between interactome and localization datasets are not directly comparable (i.e. same colors are not meant to represent the same exact set of proteins). **(C)** The hierarchical structures derived from interactome (left) and localization (right) datasets are compared to the hierarchical structures derived from “scrambled” controls – that is, to the hierarchical structure that is expected by chance given the proteins present in our dataset. Controls are generated by randomly shuffling the membership of each protein between spatial clusters or interaction communities. The number of proteins in each cluster or community was preserved from the original data.

**Figure S10:**
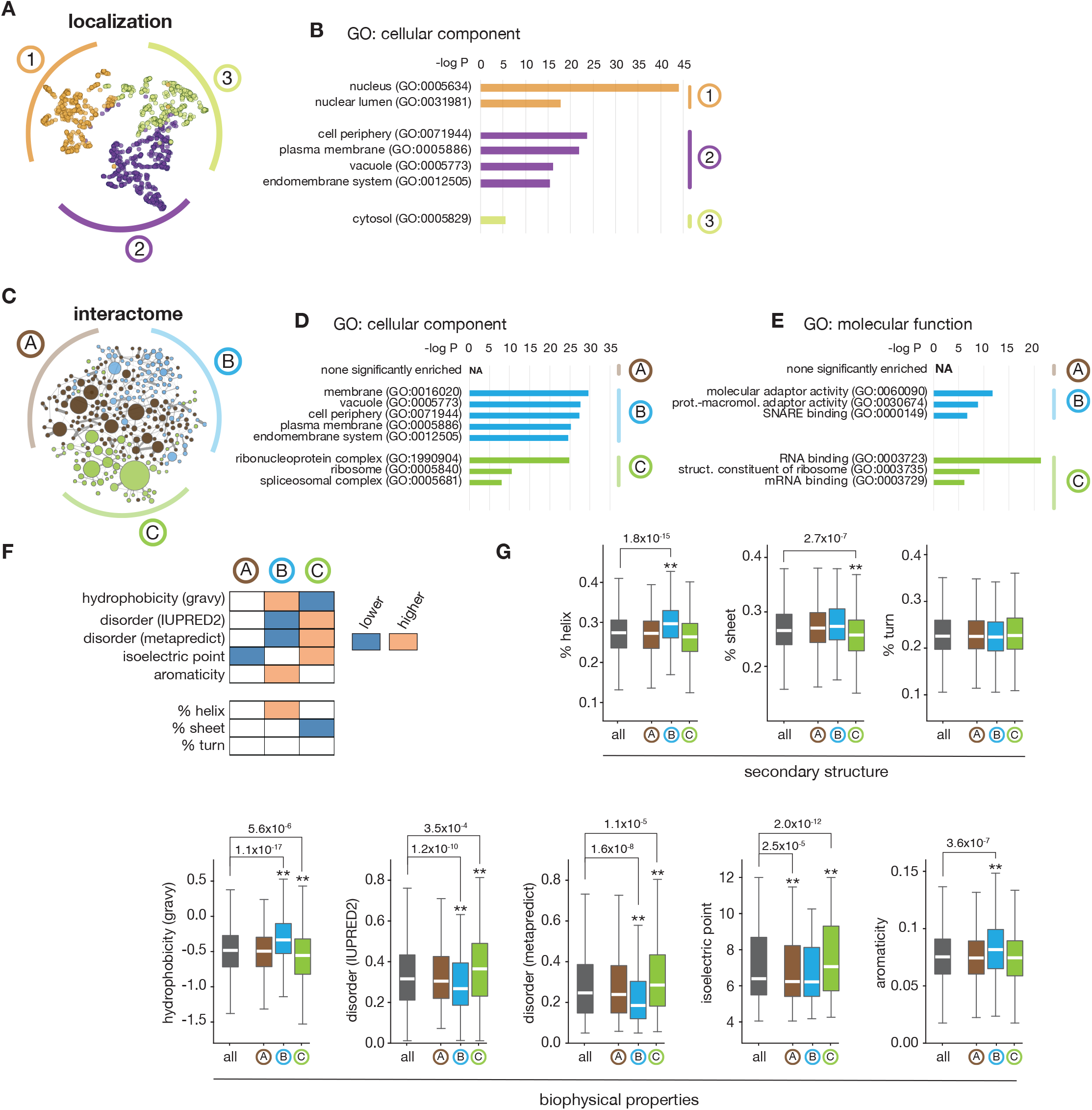
biophysical & ontology analysis of the main branches from interactome and localization hierarchies (related to Figures 5 and S9). **(A)** The three branches derived from the image-based hierarchy (see Figure S9A). **(B)** Enrichment analysis of GO annotations in the hierarchical branches, testing GO term enrichment of proteins in each branch against all proteins in the interactome (Fisher’s exact test, showing annotations enriched at p < 10^−10^ and excluding near-synonymous annotations). **(C)** The three branches derived from the interactome hierarchy (see Figure S9B). **(D)**, **(E)** Enrichment analysis of GO annotations in the hierarchical branches, testing GO term enrichment of proteins in each branch against all proteins in the interactome (Fisher’s exact test, showing annotations enriched at p < 10^−10^ and excluding near-synonymous annotations). **(F)** Heat-map representing significance testing of biophysical properties of protein sequences in the 3 branches. P-values were obtained using Student’s t-test comparing proteins belonging to a specific hierarchical branch against all proteins in the three branches. **(G)** Box plots representing the significance testing of biophysical properties described in (F). Boxes represent 25th, 50th, and 75th percentiles, and whiskers represent 1.5x inter-quartile ranges. Median is represented by a white line. ** p < 10^−3^ (Student’s t-test), exact p-values are shown.

**Figure S11:**
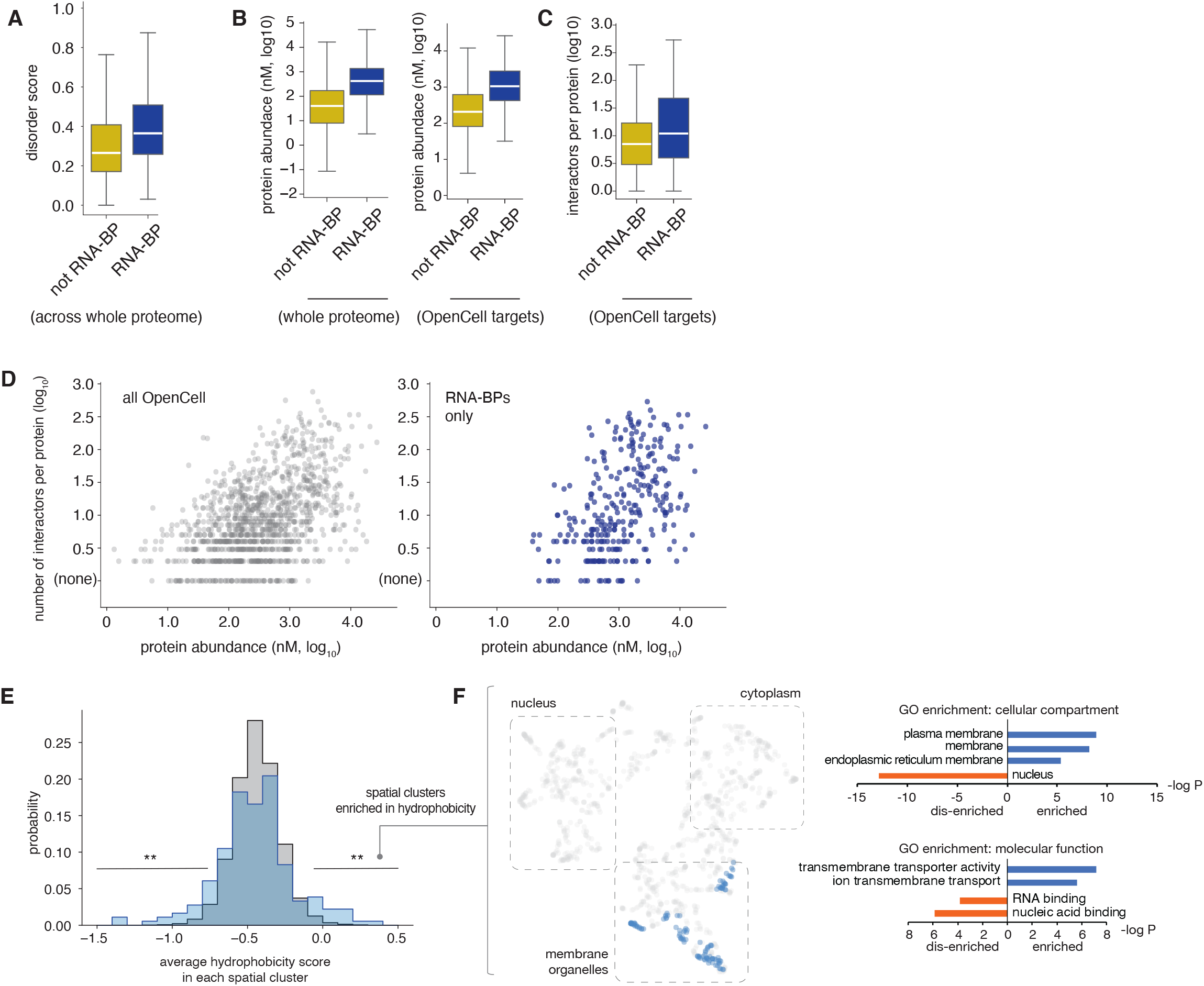
unique properties of RNA-binding proteins (RNA-BPs, related to Figure 5). **(A)** Distribution of disorder score (IUPRED2) for RNA-BPs vs. non-RNA-BPs across the whole proteome. **(B)** Distribution of protein abundance for RNA-BPs vs non-RNA-BPs across the whole proteome (left) and across OpenCell targets only (right). **(C)** Distribution of number of interactors for RNA-BPs vs non-RNA-BPs across OpenCell targets. **(D)** For each OpenCell target, the number of interactors is plotted as a function of protein abundance. The subset of targets that are RNA-BPs is highlighted on the right-hand panel. Note: for boxplots in (A), (B), (C) and (D), boxes represent 25th, 50th, and 75th percentiles, and whiskers represent 1.5x interquartile range. Median is represented by a white line. **(E)** Distribution of hydrophobicity score (gravy) across spatial clusters, comparing our data to a control in which the membership of proteins across clusters was randomized 1,000 times. Lines indicate parts of the distribution over-represented in our data vs control (**: p < 2×10^−3^, Fisher’s exact t-test). **(F)** Distribution of high-hydrophobicity spatial clusters (average hydrophobicity score > −0.1) in the UMAP embedding from Fig. 3D (left), and ontology enrichment analysis of proteins contained in these clusters (right). Enrichment compares to the whole set of OpenCell targets (p-value: Fisher’s exact test).

**Figure S12:**
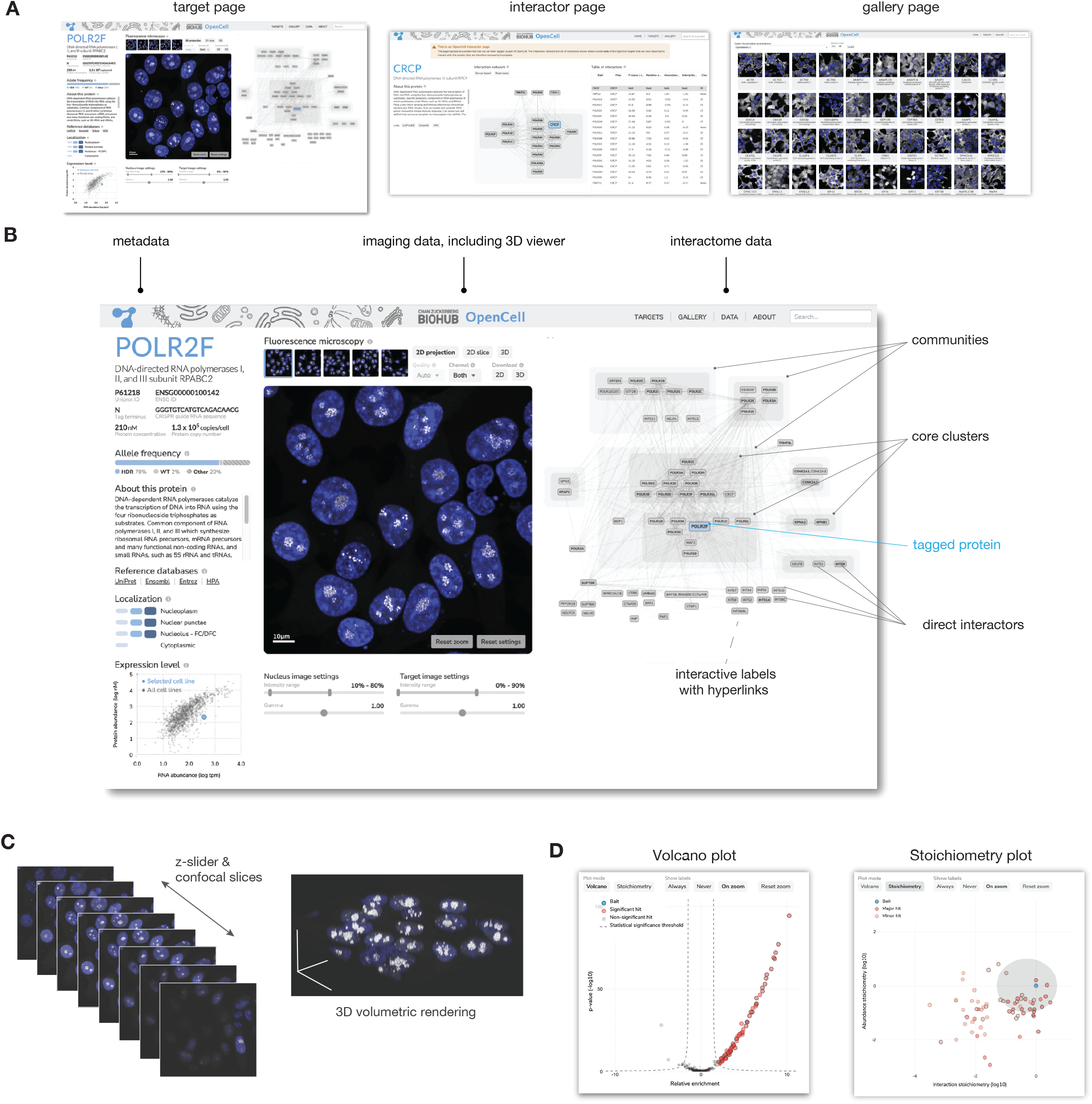
interactive data exploration at opencell.czbiohub.org. **(A)** The three principal pages of the OpenCell web app. From left to right: the target page, interactor page, and gallery page. **(B)** The target page consists of three columns. The leftmost column contains the functional annotation for the target from UniProt, links to other databases, our manually-assigned localization annotations, and measures of protein expression. The middle column contains the image viewer, and the rightmost column the interaction network. **(C)** The image viewer allows the user to scroll through the confocal z-slices using a slider or to visualize the z-stack in 3D as a volume rendering; in either mode, the user can pan and zoom by clicking, dragging, and scrolling. **(D)** The interaction network can be toggled with two alternative, complementary visualizations of the target’s protein interactions: a volcano plot of relative enrichment vs. p-value and a scatterplot of interaction stoichiometry vs. abundance stoichiometry. In both the network view and the scatterplots, the user can click on an interactor to open the target or the interactor page for the corresponding protein.

