## Supplementary material and methods for "OpenCell: proteome-scale endogenous tagging enables the cartography of human cellular organization"

[1] Chan Zuckerberg Biohub, San Francisco, USA; [2] Proteomics and Signal Transduction, Max-Planck Institute of Biochemistry, Martinsried, Germany; [3] Science for Life Laboratory, School of Engineering Sciences in Chemistry, Biotechnology and Health, KTH – Royal Institute of Technology, Stockholm, Sweden; [4] Department of Biochemistry and Biophysics, University of California, San Francisco, USA; [5] Department of Pharmaceutical Chemistry, University of California, San Francisco USA; [5] Chan Zuckerberg Initiative, Redwood City, USA; [6] Whitehead Institute, Koch Institute and Department of Biology, Massachusetts Institute of Technology, and Howard Hughes Medical Institute, Cambridge, USA; [7] Department of Cellular and Molecular Pharmacology, , University of California, San Francisco, USA; [8] NNF Center for Protein Research, Faculty of Health and Medical Sciences, University of Copenhagen, Denmark

### <u>Supplementary materials</u>

#### Materials and Methods

##### Cell culture and CRISPR engineering

**Cell culture.** HEK-293T cells (ATCC CRL-3216) were cultured in DMEM high-glucose medium (Gibco, cat. #11965118) with 10% fetal bovine serum (Omega Scientific, cat. #FB-11), supplemented with 2mM glutamine (Gibco, cat. #25030081), penicillin and streptomycin (Gibco, cat. #15140163). All cell lines were maintained at 37°C and 5% CO2 and routinely tested for the absence of mycoplasma.

**Fluorescent library design.** mNeonGreen is monomeric green fluorescent protein ~2x brighter than GFP. We used the split-mNeonGreen$_2$ system for functional tagging, which separates last mNeonGreen$_2$ beta-strand (mNG11) from the rest of the fluorescent protein (mNG1-10)(*20*). Upon co-expression in the same cell, mNG1–10 and mNG11 stably assemble and reconstitute a functional FP. A parental cell line constitutively expressing mNG1-10 was first generated by lentiviral transduction (from pSFFV_mNG$_2$1-10, Addgene #82610). All successive cell lines were generated from this parental HEK293T$^{mNG1-10}$ cell line by incorporation of the mNG11 fragment at either N- or C- terminus of the genomic sequence of a target protein via CRISPR/Cas9 based genome editing. Our mNG11 fusion constructs include a HRV 3C cleavable linker(*91*) that can be used optionally for elution from an affinity capture matrix (16 a.a. tag + 14 a.a. linker, full sequences in Table S1). To minimize the risk of functional perturbation, we stringently selected integration sites (N- or C-terminus) by systematically curating the literature for data supporting the functional integrity of fusion proteins (or by requesting advice from cell biology experts for specific proteins). We also used 3D PDB structures whenever available to identify sites that avoid protein-protein interaction interfaces. See Table S1 for details. Because our split-FP system does not enable detection in the lumen of organelles (this requires split constructs harboring appropriate signal sequences(*23*)), fusions with membrane proteins were restricted to cytoplasmic termini, ensuring first that no annotated regulatory sequences (e.g., signal sequences) were compromised. In total, we used available supporting data to inform 62 % of insertion sites, and 3 % were constrained by membrane protein topology. In the absence of prior information, insertion choice was based on avoiding annotated regulatory sites. In the case of splice variants involving the terminus of choice, the main transcript expressed in HEK293T was used.

**Overall genome engineering pipeline.** To enable the expression of fluorescent fusion from endogenous genomic loci, we used an established high-throughput CRISPR/Cas9 method for gene editing by homologous recombination (*18*). In brief, *S. pyogenes* Cas9/guide RNA complexes were pre-assembled in vitro, mixed with short single-stranded oligo-nucleotide homology donors and delivered into HEK293T$^{mNG1-10}$ cells by electroporation in 96-well plates (see below). For each genomic insertion, the choice of guide RNA and associated homology donor sequence (which contains the mNG11 payload flanked by short sequences of genomic homology to the targeted insertion site) was automated using *crispycrunch*(*92*), an open-source CRISPR design software available at github.com/czbiohub/crispycrunch and as a web-app at crispycrunch.czbiohub.org/. *crispycrunch* selects a guide RNA closest to a desired genomic insertion site while also minimizing any off-target guide RNA activity, and if needed introduces silent mutations to inactivate guide

RNA biding and re-cutting after successful homologous recombination (*92*). gRNA and homology donor sequences for all targets are found in Table S1.

**Cell engineering and selection** *S. pyogenes* Cas9 protein (pMJ915 construct, containing two nuclear localization sequences) was expressed in E. coli and purified by the UC Berkeley Macrolab following protocols described by Jinek et al(*93*). Cells were synchronized by nocodazole treatment (200ng/mL for 15-18h_ to enhance homologous recombination(*25*). RNP complexes were freshly assembled with 50 pmol Cas9 protein and 65 pmol gRNA prior to electroporation, and combined with HDR template in a final volume of 10 µL. First, 0.5 µL gRNA (130 µM stock) was added to 2.35 µL high-salt RNP buffer {580 mM KCl, 40 mM Tris-HCl pH 7.5, 20% v/v glycerol, 2 mM TCEP-HCl pH 7.5, 2 mM MgCl2, RNAse-free} and incubated at 70°C for 5 min. 1.25 µL of Cas9 protein (40 µM stock in Cas9 buffer, ie. 50 pmol) was then added and RNP assembly carried out at 37°C for 10 min. Finally, HDR templates and sterile RNAse-free H2O were added to 10 µL final volume. Electroporation was carried out in Amaxa 96-well shuttle Nucleofector device (Lonza) using SF solution (Lonza) following the manufacturer's instructions. Cells were washed with PBS and resuspended to 10,000 cells/µL in SF solution (+ supplement) immediately prior to electroporation. For each sample, 20 µL of cells (ie. 200,000 cells) were added to the 10 µL RNP/template mixture. Cells were immediately electroporated using the CM130 program, after which 100µL of pre-warmed media was added to each well of the electroporation plate to facilitate the transfer of 25,000 cells to a new 96-well culture plate containing 150µL of pre-warmed media. Electroporated cells were cultured for >5 days and transferred to 12-well plates prior to selection by fluorescence-activated cell sorting (FACS). For each target, 1,200 cells from the top 1% fluorescent cell pool were isolated on a SH800 instrument (Sony biotechnology) and collected in 96-well plates.

**Genotype analysis.** For each polyclonal pool of engineered cells, the genotype of CRISPR-edited alleles was characterized by amplicon sequencing. Gene-specific primers were designed using Primer3, with a target amplicon length of 270bp and a maximum at 500bp. gDNA was first extracted by cell lysis using QuickExtract DNA Extraction Solution (Lucigen). From a confluent culture in 96-well plate, media was removed, cells were washed 1x in DPBS and resuspended in 50 µL QuickExtract. The cell layer was detached by repeated pipetting and transferred to a PCR plate for incubation. The lysate was incubated as follows {65°C for 20 min, 98°C for 5min, 4°C final}. gDNA was used directly from this preparation. Amplicon Libraries were created using a two-step PCR protocol: the first PCR amplifies the target genomic locus and adds universal amplification handle sequences, while the second PCR introduces index barcodes using the universal handles. PCR1: this PCR uses a "reverse touchdown" method designed to accommodate a number of different annealing temperatures for a number of different targets. 50-µL PCR reactions were set using 2x KAPA HiFi Hotstart reagents (Roche) with 2µL extracted gDNA, 80pmol each primer and betaine to 0.8M final concentration. PCR conditions: 95°C 3min; 3 cycles of {98°C for 20s, 63°C for 15s, 72°C for 20s}, 3 cycles of {98°C for 20s, 65°C for 15s, 72°C for 20s}, 3 cycles of {98°C for 20s, 67°C for 15s, 72°C for 20s}; 17 cycles {98°C for 20 s, 69°C for 15 s, 72°C for 20s} then 72°C for 1min; 4°C final. PCR2: amplicons were diluted 1:100 and 1 µL was used into a 40-µL barcoding reaction using 20 µL 2x KAPA HiFi Hotstart reagents (Roche) and 80pmol each barcoded primer. PCR conditions: 95°C 3min and 12 cycles of {98°C for 20s, 68°C for 15s, 72°C for 12s} then 72°C for 1min; 4°C final. Barcoded amplicons were analyzed using capillary electrophoresis (Fragment Analyzer, Agilent), pooled and purified using magnetic beads. Sequencing was performed on an Illumina Miseq V3 platform (paired-end 2x300bp) using standard P5/P7 primers. Genotype analysis was performed using CRISPRESSO2, which allowed to quantify three classes of alleles for each targeted locus: un-modified (wild-type), alleles integrated with mNG11 by homologous recombination, and alleles containing non-functional mutations as a result of competing DNA repair mechanisms. Primer sequences and genotype analysis for all targets are found in Table S1. Despite multiple attempts, genotyping PCR could not be successfully performed for 70 targets (5% of the total set), most often involving genes with extreme GC content or highly repetitive sequences.

#### Immuno-precipitation / mass-spectrometry

**Overall strategy.** mNG11-tagged proteins were isolated from digitonin-solubilized lysates using anti-mNeonGreen nanobody capture. Triplicate protein samples were digested "on-bead" for bottom-up proteomics analysis(*28*), and peptides were quantified using label-free mass spectrometry on a timsTOF Pro instrument (Bruker Daltonics).

**Sample preparation.** Confluent 12-well cultures (0.8x$10^6$ cells/sample) were washed twice with 1 ml of D-PBS (no divalent). 200 µl ice-cold lysis buffer A {50 mM HEPES pH 7.5, 150 mM KOAc, 5 mM NaCl, 2 mM MgOAc, 1 mM $CaCl_2$, 15% Glycerol, 1.5 % Digitonin (high purity, Calbiochem), Protease- and Phosphatase inhibitor (Halt, Pierce), 0.1% benzonase (Millipore Sigma)} were added to each well, cells were lysed by strong pipetting and the solution was transferred into a pre-chilled 96-well PCR plate. Per 96-well plate, 330 µl magnetic mNG-Trap slurry (magnetic agarose, Chromotek) was washed three times with buffer B {50 mM HEPES pH 7.5, 150 mM KOAc, 5 mM NaCl, 2 mM MgOAc, 1 mM $CaCl_2$, 15% Glycerol, 0.1 % Digitonin} and resuspended in 2,150 µl Buffer A. The cell lysate was incubated for 1h at 4°C, rotating. The insoluble cell fraction was pelleted for 30 min at 1800xg in a table-top centrifuge at 4°C, followed by supernatant transfer into a new plate pre-loaded with 20 µl of the washed bead slurry per well. Tagged proteins were captured by incubation for 2h at 4°C, rotating. Following capture and using a 96-well magnet, beads were washed (per well) with 200 µl buffer B (incubation for 5 min at 4°C, rotating), 2x 200 µl buffer B (no incubation) and a final 1x 200 µl buffer C to remove digitonin {50 mM HEPES pH 7.5, 150 mM KOAc, 5 mM NaCl, 15% Glycerol, 0.01% glyco-diosgenin (Avanti)}. Supernatant was removed and 50 µl of digestion buffer 1 {6 M Urea, 50 mM Tris-HCl, pH 8.5, 1 mM DTT, 2 ng/µl LysC protease (Wako Chemicals)} was added to each well, followed by overnight digestion at 30°C on a thermomixer, gently shaking. The next day, 100 µl digestion buffer 2 {50 mM Tris-HCl, pH 8.5, 8.25 mM iodoacetamide, 2 ng/µl LysC} was added to each well and incubated for ~6 hours at 30°C on a thermomixer in the dark, gently shaking. The digestion was finally quenched with 15 µl of 10 % TFA. Quenched samples were vortexed, flash-frozen and stored at -80 °C until further use for LC-MS analysis preparation.

**EvoSep chromatography.** We used the EvoSep liquid chromatography system for sample processing(*94*). EvoTips (EvoSep Gmbh) were activated for 5 min with 1-Propanol at RT, followed by a wash step with 50 µl Buffer A (99.9 % ddH2O, 0.1 % Formic Acid) and centrifugation at 600 xg for 1 min at RT. The flow-through was discarded and activated EvoTips were placed in an EvoTip-box reservoir filled with Buffer A. After on-bead digestion, captured protein samples were thawed for 5 min at 600 rpm and 25°C on a thermal shaker and placed on a 96-well magnet holder to remove magnetic beads. The whole sample (~150 µl) was transferred to activated EvoTips, followed by two consecutive centrifugation steps at 600xg for 1 min and RT, discarding flow-through after the first spin. Peptide-loaded EvoTips were washed once with 50 µl Buffer A and centrifuged at 600xg for 1 min at RT. The flow-through was discarded and 150 µl of Buffer A was added to each EvoTip followed by a centrifugation step for 20 sec at 600xg RT. Loaded EvoTips were then transferred into the 96-well EvoTip-box reservoir filled with Buffer A and transferred onto the EvoSep autosampler for LC-MS analysis. Pulldowns were acquired in triplicates and injected to the mass spectrometer while spacing replicates to prevent any bias.

**Liquid-chromatography.** For separating peptides by hydrophobicity and eluting them into the mass spectrometer, we used an EvoSep One1 liquid chromatography system (EvoSep, Gmbh) and analyzed purified petides with a standard 21 min method (60 samples per day). We used a 15 cm × 150 µm ID column with 1.9 µm C18 beads (PepSep) coupled to a 20 µm ID electrospray emitter (Bruker Daltonics). Mobile phases A and B were 0.1 % FA in water and 0.1 % FA in ACN, respectively. The EvoSep system was coupled online to a trapped ion mobility spectrometry quadrupole time-of-flight mass spectrometer(*95*) (timsTOF Pro, Bruker Daltonics) equipped with via a Captive nano-electrospray ion source.

**Mass spectrometry.** Mass spectrometric analysis was performed in a data-dependent (dda) PASEF mode. For ddaPASEF, 1 MS1 survey TIMS-MS and 4 PASEF MS/MS scans were acquired per acquisition cycle. The cycle overlap for precursor scheduling was set to 2. Ion accumulation and ramp time in the dual TIMS analyzer was set to 50 ms each and we analyzed the ion mobility range from 1/K0 = 1.3 Vs cm-2 to 0.8 Vs cm-2. Precursor ions for MS/MS analysis were isolated with a 2 Th window for m/z < 700 and 3 Th for m/z >700 in a total m/z range of 100-1,700 by synchronizing quadrupole switching events with the precursor elution profile from the TIMS device. The collision energy was lowered linearly as a function of increasing mobility starting from 59 eV at 1/K0 = 1.6 VS cm-2 to 20 eV at 1/K0 = 0.6 Vs cm-2. Singly charged precursor ions were excluded with a polygon filter (otof control, Bruker Daltonics). Precursors for MS/MS were picked at an intensity threshold of 2,000 arbitrary units (a.u.) and re-sequenced until reaching a 'target value' of 24,000 a.u. considering a dynamic exclusion of 40 s elution. Capillary voltage was set to 1,750 V and dry gas temperature to 180°C.

**Raw Data Processing.** MS raw files were processed using MaxQuant (v1.6.10.43)(*96*, *97*), which extracts features from four-dimensional isotope patterns and associated MS/MS spectra, on a computing cluster (SUSE Linux Enterprise Server 15 SP2) utilizing UltraQuant. Files were processed in several batches of appriximately 1000 files each and searched against the human Uniprot databases (UP000005640_9606.fa, UP000005640_9606_additional.fa). False-discovery rates were controlled at 1% both on peptide spectral match (PSM) and protein levels. Peptides with a minimum length of seven amino acids were considered for the search including N-terminal acetylation and methionine oxidation as variable modifications and cysteine carbamido-methylation as fixed modification, while limiting the maximum peptide mass to 4,600 Da. Enzyme specificity was set to LysC cleaving c-terminal to lysine. A maximum of two missed cleavages were allowed. Maximum precursor and fragment ion mass tolerance were searched as default for TIMS-DDA data and the main search tolerance was reduced to 20 ppm. Peptide identifications by MS/MS were transferred by matching four-dimensional isotope patterns between the runs (MBR) with a 0.7-min retention-time match window and a 0.05 1/K0 ion mobility window. Protein quantification was performed by label-free quantification using a minimum ratio count of 1.

**Data availability.** All mass spectrometry raw data and MaxQuant output tables are deposited to the ProteomeXchange Consortium(*98*) via the PRIDEpartner repository and will be publicly available upon final publication (accession PXD024909).

#### Whole-cell abundance measurement by mass-spectrometry

**Peptide preparation.** HEK293T cells were grown in triplicate 15cm-plates, washed 2x in ice-cold PBS and lysed in { 2.5% SDS sodium dodecyl-sulfate; 50 mM Tris pH 8.1 }. Lysis was performed at 95˚C for 5 min, followed by probe sonication. Lysates were cleared by centrifugation, protein amount was measured by BCA assay, and lysates were precipitated with 5 volumes of acetone. Pellets were resuspended in 50 mM Tris pH 8.1 containing 8 M urea, reduced with 1 mM DTT and alkylated with 5 mM IAA before initiation of digestion overnight with LysC at an enzyme-to-protein ratio of 1:100. The digest mixture was diluted four-fold, and trypsin was added at an enzyme-to-protein ratio of 1:100 for 6 h, followed by an additional aliquot of trypsin overnight. The digestion reaction was stopped by acidifying the sample adding TFA to 1%, placed on ice for 10min and centrifuged at 4 degree C, 21000g for 20min. The resulting peptide supernatant was then desalted using mixed mode Strata-XC SPE cartridge. Briefly, the cartridge was prepared by activating with methanol, conditioning with 80% acetonitrile/0.1% TFA and equilibrated with 0.2% TFA. The acidified peptides were then added, washed with 99% isopropanol/0.1% TFA, 2 x 0.2% TFA washes, 1x 0.1% formic acid and eluted with 60% acetonitrile/0.5% ammonium hydroxide. The eluted peptides were flash frozen and then dried down.

**Fractionation.** To obtain achieve measurement depth, peptides from the triplicate experiment were further separated in 24 fractions using C18 chromatography. Peptides were resuspended in buffer A (10 mM ammonium bicarbonate) and injected onto a 4.6 × 250-mm 3.5-μm Zorbax 300 Extend-C18 column. Peptides were separated on a non-linear gradient as described in (*99*), using the following composition of buffer B (10 mM ammonium bicarbonate, 90% acetonitrile). Peptide fractions were frozen at −80 °C before centrifugal evaporation.

**Mass spectrometry.** Peptides were resuspended in 2% ACN with 0.1% TFA before loading onto a 25 cm x 75 μm ID, 1.6 μm C18 column (IonOpticks) maintained at 40°C. Peptides were separated with an EASY-nLC 1200 system (Thermo Fisher Scientific, San Jose, CA) at a flow rate of 300 nl min-1 using a binary buffer system of 0.1% FA (buffer A) and 80% acetonitrile with 0.1% FA (buffer B) in a two-step gradient, from 3% to 27% B in 105 min and from 27% to 40% B in 15min. All fractions were analyzed on a Fusion Lumos mass spectrometer (Thermo Fisher Scientific, San Jose, CA) equipped with a nanoFlex ESI source operated at 1550 volts, RF lens set to 30%, operated in data dependent acquisition mode with a duty cycle time of 1 sec. Full MS scans were acquired with a m/z scan range of 375-1500 m/z in the Orbitrap mass analyzer (FTMS) with a resolution of 240k. Selected precursor ions were subjected to fragmentation using higher-energy collisional dissociation (HCD) with a quadrupole isolation window of 0.7 m/z, and normalized collision energy of 31%. HCD fragments were analyzed in the Ion Trap mass analyzer (ITMS) set to Turbo scan rate. Fragmented ions were dynamically excluded from further selection for a period of 45 sec. The AGC target was set to 1,000,000 and 10,000 for full FTMS and ITMS scans, respectively. The maximum injection time was set to Auto for both full FTMS and ITMS scans.

#### Live-cell imaging

**Sample preparation.** Live-cell imaging was performed on 96-well glass-bottom plates (Greiner Bio One, cat. #655891) coated with 50µg/ml fibronectin (Corning, cat. #356008). Cells were seeded on an imaging plate 28-32 hours before imaging at 15,000 cells per. Before imaging, cells were counter-stained with the live-cell DNA dye Hoechst 33342 (Invitrogen, cat. #H3570) by incubation for 30 minutes at 37°C in 150 µl of Hoechst diluted to 1µg/mL in culture media. Media was then replaced with phenol-free DMEM (Gibco, cat. #21063029) supplemented with 10% FBS. Hoechst staining was performed three to four hours prior to imaging to provide the cells time to recover from any mechanical stress due to medium changes.

**Live-cell fluorescence microscopy.** Cells were imaged on a DMI-8 inverted microscope (Leica) equipped with a Dragonfly spinning-disk confocal system (Andor), a 63x 1.47NA oil objective (Leica), and a 16-bit iXon Ultra 888 EMCCD camera (Andor, pixel size: 13x13 $\mu m^2$). A pinhole size of 40µm was used with an EM gain of 400. Cells were maintained at 37°C and 5% CO2 during image acquisition by a stage-top incubator (Okolab, H101-K-Frame). The microscope was controlled using the open-source microscope-control software MicroManager (version 1.4.22).

**Automated confocal acquisition**. We automated the imaging of 96-well plates using a custom acquisition script, written in Python, combined with a custom MicroManager plugin (mm2python; github.com/czbiohub/mm2python) to expose the MicroManager APIs in a Python environment. This script selected optimal fields of view (FOVs) at which to acquire confocal z-stacks by using a pre-trained machine-learning model to assign a quality score to the FOVs at a set of different positions in each well. Briefly, at each position, the script acquired a single 2D snapshot of the Hoechst staining, segmented the nuclei in the snapshot, and calculated an array of features associated with the distribution of nuclei within the FOV. The script then used a pre-trained random-forest regression model (see below) to predict a quality score for the FOV from this set of features. This process was repeated at each of 25 different positions in each well, and then the script selected the positions with the highest-scoring FOVs to revisit for confocal z-stack acquisition. At each of these selected positions, the focal plane was centered on the cell layer using a laser-based Adaptive Focus Control system (Leica) and confocal z-stacks, consisting of 110 z-slices at a spacing of 0.2µm, were acquired. The exposure settings for the mNeonGreen channel were determined dynamically for each target using a custom auto-exposure algorithm that iteratively adjusted the exposure time and laser power until the maximum pixel intensity was just below or just above an intensity of $2^{15}$ (half of the full dynamic range of the camera). For dim targets for which this condition could not be met, the script fell back to a hard-coded absolute maximum exposure time and laser power to minimize both acquisition time and photobleaching. The exposure settings for the Hoechst stain were manually selected and held constant for all targets. The random-forest regression model used by the script to predict the FOV quality scores was trained prior to acquisition using a set of 3800 FOV snapshots that were manually assigned to one of three grades: “poor,” “mediocre,” or “good.” These grades were mapped to a continuous response variable by assigning the values of -1, 0, and 1, respectively, and a random forest regression model (scikit-learn) was trained to predict this value. The out-of-bag estimated $R^2$ was 0.86 and scores predicted for a withheld set of test snapshots were also evaluated by manual inspection. The trained model was cached and imported at acquisition time by the acquisition script. The acquisition script, trained FOV-scoring model, autoexposure algorithm, and other associated microscope-control methods are available online at github.com/czbiohub/2021-opencell-microscopy-automation.

#### Data analysis – proteomics

**Statistical detection of protein interactions.** Statistical analysis was performed according to methods described in Hein et al. (*7*), with modifications. Protein identifications were filtered, removing common contaminants, hits to the reverse decoy database as well as proteins only identified by modified peptides. We required that each protein be quantified in all replicates from the IP-MS samples of at least one cell line and used log2 MaxQuant LFQ intensities for all analyses. Rather than imputing missing values, robust null control sets were generated for statistical enrichment analysis of each protein group by pooling triplicate data from an average of 349 unrelated samples. In this approach, rather than using a single control we measure enrichment in a specific sample against an entire cohort of ~349 unrelated tagged cell lines. We have previously described (*7*, *28*) how this enables a better estimation of the null distribution and leads to more robust identification of interactions. The null control sets might contain triplicate samples that are outliers and would be considered significant interactions. The presence of these samples lead to underestimation of enrichment and could mask some significant interactions. We systematically removed these outliers from the negative control sets using a Student's t-test and excluding any sample of triplicates that had a p-value < 0.001. From the filtered pool, we approximated the true mean and the true standard deviation of the null set by bootstrapping via sampling with replacement. The approximated mean and standard deviation of the null set was then used for the final Student's t-test to calculate the statistical significance of the triplicate means. Any missing values in the triplicate sample set were then replaced with the mean of the null set. Enrichment was calculated by subtracting the mean of the triplicates from the mean of the null set, and was normalized to account for variability within each protein through division by the standard deviation of the null control set. Our statistical strategy to define significant interactors is described in Figure S4A-B and supported by a quantitative estimation of precision and recall.

**Precision / recall analysis of the interactome.** For a quantitative evaluation of our statistical approach, and to compare the quality of OpenCell against reference interactome datasets, we created a framework the precision and recall in interaction data. In the absence of established ground truth for human protein interactions, we indirectly derived measurements of precision and recall. For recall, we calculated the coverage in a given dataset of interactions curated in the human CORUM database(*30*), as a percentage of all possible CORUM interactions given the set of baits in that dataset. For calculating precision, we used the assumption that two interactors should have localization patterns that at least partially overlap. As an independent ground truth set for protein localization, we used the quantitative analysis of the HeLa proteome from Itzhak et al. (*31*). Using these annotations, we categorized localization into four broad classes: exclusively nuclear, exclusively cytoplasmic, exclusively organellar, and multi-localizing (i.e., any non-exclusive localization). To calculate precision, we consider any two interactors that overlap in exclusive localization to be true positives, and those that do not overlap localization annotations at all to be false positives, with multi-localizing proteins allowed to interact agnostically (Fig. S4B).

**Protein stoichiometry measurements.** Calculation of interaction stoichiometries was performed as in Hein et al by dividing LFQ intensities by the number of theoretically observable peptides for each protein. We defined the "interaction stoichiometry" as the stoichiometry of the abundance of a given interactor, relative to the abundance of the corresponding bait, in a given pull-down. We also defined a "cellular abundance stoichiometry" as the stoichiometry of the abundance of a given interactor, relative to the abundance of the corresponding bait, in a whole cell lysate. For proteins that were not detected in whole cell lysates (due to lack of measurable peptides, for example in the absence of lysine residues), protein abundances were imputed from RNA-Seq data by interpolating from a linear regression of RNA-Seq tpm vs. protein abundance measured by mass spectrometry in our dataset.

**Network Analysis.** For graph-based clustering of the entire interactome network, we weighted edges using the interaction stoichiometry between each pair of interacting proteins. We utilized Markov clustering(*34*) at various inflation parameters and evaluated clustering performance using the k-clique method described in Drew et al(*100*) using CORUM complexes as the ground truth. To eliminate complexes with many shared proteins, the Jaccard distance was calculated between all pairs of complexes, and pairs of complexes were merged if the distance was below 0.6. Our final clustering analysis used an inflation parameter of 3.0 (Fig. S4I). The clusters were pruned to remove any node included in a cluster on the basis of a single edge. The resulting clusters correspond to the protein "communities" described in the text. We then utilized another round of MCL clustering to identify core-clusters within each community by considering only highly stoichiometric interactions (interaction stoichiometries between 0.05 and 10, and cellular abundance stoichiometry between 0.1 and 10). The resulting core-clusters represented highly stable core clusters within the original communities.

**Measurement of biophysical properties of proteins.** Biophysical properties were calculated using the *ProteinAnalysis* package from BioPython(*101*). Hydrophobicity scores were calculated using the *gravy* method of that module to compute the Gravy index. Calculation of disorder in protein regions was performed using the IUPred2A algorithm(*89*) or metapredict, a recent agglomerative algorithm (*90*). Scores were averaged across the sequences of each protein. Scores computed across the whole proteome are included in Table S2.

#### Data analysis – imaging

**Consensus localization encodings.** Protein localization patterns were encoded from the raw confocal images using a customized variant of the vector-quantized autoencoder architecture VQ-VAE-2(*102*). The image preprocessing, autoencoder architecture, and model training are described in detail in an accompanying manuscript(*52*). Briefly, confocal z-stacks were reduced to two dimensions by a maximum-intensity z-projection and normalized to control for variation in intensity. Regions of interest 200x200 pixels in size were centered on individual nuclei and cropped from each z-projection to generate a set of 50-200 cropped images for each tagged protein. These images were randomly partitioned into a training set and a test set. After training the model on the images in the training set, the images in the test set were encoded, and the resulting latent-space vectors from the VQ2 layer of the network were flattened to obtain a localization encoding for each image in the test set as a 9216-dimensional vector. The encodings of all images for each tagged protein were then averaged to obtain a single consensus encoding for each tagged protein. The matrix of consensus encodings for all OpenCell targets are available on Figshare at:

https://figshare.com/articles/dataset/Consensus_protein_localization_encodings_for_all_OpenCell_targets/16754965

**Comparison of OpenCell and Human Protein Atlas (HPA) localization annotations.** The v20 dataset of HPA localization annotations was first obtained from the HPA website (https://v20.proteinatlas.org/download/subcellular_location.tsv.zip). To compare OpenCell localization annotations to HPA annotations, it was necessary to reconcile the OpenCell and HPA localization categories as the ontologies used to annotate the two datasets vary slightly. To do so, a set of 'consensus' annotation categories were defined for the most common localization categories, as described in Table S7 (see sheet: "annotation-definitions").

Because wide-spread multi-localization of proteins complicates direct comparisons, we focused first on comparing the "main" localization annotations provided by each dataset. After mapping to these consensus categories, grade-2 and grade-3 OpenCell annotations were compared to their corresponding HPA 'main location' annotations and categorized as either exact matches, partial matches, or entirely discrepant. Exact matches were targets whose sets of consensus annotations were identical in the OpenCell and HPA datasets; partial matches were targets with at least one of the same consensus annotations in the OpenCell and HPA datasets. The list of exact and partial matches, and the sets of consensus OpenCell and HPA annotations, are provided in Table S7.

To refine the list of proteins that did not share any matching annotation across the datasets, minor localization annotations were considered (grade 1 in OpenCell, "additional" localization in HPA), as well localization between closely related organelles (for example, ER and Golgi), which could explain differences between the datasets as they probe localization in different cell lines. As a result, a final list of 147 proteins for which the two dataset were fully discrepant was obtained. The full analysis of proteins from that list using literature curation is presented in Table S8.

**Analysis of image localization encodings.** The matrix of consensus localization encodings for all OpenCell targets was analyzed using the *scanpy* package(*103*). Briefly, the dimensionality of the consensus encodings was reduced using PCA and the first 200 PCs, which captured 96% of the variance, were retained for downstream analysis. The UMAP algorithm (*53*) was used to embed the encodings in two-dimensional space using 10 nearest neighbors, the Euclidean distance metric, and a minimum embedding distance of zero. The encodings were clustered using the Leiden graph-based clustering algorithm(*55*) with a resolution parameter of 30 and the weighted adjacency matrix calculated by the UMAP algorithm (again with 10 nearest neighbors). Finally, the Pearson correlation coefficient between the top 200 PCs of the localization encodings was used to quantify the localization similarity between OpenCell targets.

**Image-based clustering.** OpenCell targets were clustered on the basis of their consensus encodings using the Leiden graph-based clustering algorithm (*55*) and the weighted adjacency matrix calculated by the UMAP algorithm with 10 nearest neighbors. The Leiden algorithm depends upon a single 'resolution' hyperparameter that determines the number of clusters. To quantify clustering performance as a function of this hyperparameter, the Adjusted Rand Index (*88*) was used to compare the Leiden clusters to ground-truth datasets. The ARI is near zero for random clustering and is equal to one when clustering perfectly matches the ground-truth labels. Three different ground-truth datasets were used that capture biological relationships at three different scales: manual OpenCell localization annotations (organelle scale), KEGG pathways (https://www.genome.jp/kegg/ (*104*)), and CORUM complexes (http://mips.helmholtz-muenchen.de/corum/ (*30*). OpenCell targets that were in more than one ground-truth cluster were excluded from this analysis, as the ARI is defined only for hard clustering (that is, sample-cluster assignments that are one-to-one). The ARI was calculated with respect to each of the ground-truth datasets at a range of values of the Leiden resolution hyperparameter; the global maxima in the resulting ARI curves correspond to the clustering resolutions that best capture the information in each ground-truth dataset. To control for the stochasticity of the Leiden algorithm, the ARI curve was calculated from the average of the curves for nine random seeds.

#### Hierachical analysis of interactions and localization patterns.

**Hierarchical clustering of interactome and image-localization clusters**. To explore the relationships between the 182 localization clusters or the 300 interactome communities, we employed the Paris algorithm, an agglomerative graph-based hierarchical clustering algorithm(*105*). The algorithm was initialized with a network of nodes representing the initial clusters (either the localization clusters or the interactome communities) and edge weights between the initial clusters were calculated according to the definition of the cluster pair sampling ratio used in the Paris algorithm.

**Gene Ontology enrichment analysis**. To analyze enrichment of GO terms in a given hierarchical protein group, we utilized the PANTHER gene list analysis API (*106*) using Fisher exact test for significance testing. Enrichment of GO terms was tested against a reference set of either all OpenCell targets for the imaging dataset, or all proteins found in communities for the interactome dataset.

#### OpenCell web portal development

The OpenCell web portal is a full-stack web application. The frontend (that is, the web interface itself) is written with React, a modern JavaScript library for building modular user interfaces. The backend is a PostgreSQL database paired with a REST API written in Python using Flask and SQLAlchemy. Together, the database and API provide the metadata, protein interaction data, and the confocal image data required to populate the frontend. For efficiency, the 3D confocal stacks are transferred to the client as two-dimensional tiled arrays of confocal slices, saved as compressed JPEG images to enable fast download times. To maximize responsiveness, the web app makes API requests dynamically and asynchronously so that it loads, in parallel, only the data required to update the state of the app in response to a given user input. Both the backend and frontend are built using many open-source packages. In particular, the 3D rendering of confocal stacks relies on Three.js, the interactive scatterplots are built with d3.js, and the interaction networks are built with Cytoscape.js. The backend is built with SQLAlchemy and Flask and also leverages the Python data-science stack, including pandas, NumPy, SciPy, and scikit-image. All source code for the application is available on GitHub at github.com/czbiohub/opencell-portal-pub

**Figure generation**

Data analysis was performed in Python. Figures were generated in Python using matplotlib or seaborn, with the exception of the protein-protein interactions / network visualizations, which were generated using Cytoscape. The code and data used to generate the figures can be found on GitHub at github.com/czbiohub/2021-opencell-figures.

**Fig. S1: Experimental pipeline (related to Fig. 1). (A)** IP-MS using FP capture. All mNG11 tagging constructs also include an HRV-3C cleavable linker for optional release from the capture resin. **(B)** Justifying the choice of tag insertion in engineered cell lines. To inform tag insertion sites, we used a combination of existing data from the literature suggesting preservation of properties, 3D structures of protein complexes from the PDB and sequence analysis to avoid important functional motifs. 4% of insertion sites were constrained by the topology of transmembrane protein targets (fusion to cytosolic termini), and for 23% of targets no prior data was available. See details in Table S1. **(C)** Sensitivity of interaction proteomics detection on a timsTOF instrument. The number of interactors detected in pull-downs from 6 different targets is shown, varying the amount of input material. To balance sensitivity and scalability, 0.8e6 cells were used for high-throughput assays (12 well-plate, wp). **(D)** Distribution of gene ontology annotations in the OpenCell library (successful targets) compared the whole proteome. Over- and under-represented terms are outlined. Because organellar organization and transport between organelles are foundational to human cellular architecture, proteins in these groups are slightly enriched in our library. Under-represented groups are mostly comprised of proteins in compartments that are not accessible to our tagging strategy (mitochondrial functions, extracellular matrix) or proteins that are typically present at low copy numbers and therefore difficult to detect at endogenous levels (transcription factors).

**Fig. S2: Cell line generation (related to Fig. 1). (A)** Success rate for the generation and detection by imaging of fluorescently tagged cell lines are compared for the whole set of targets we attempted, and the subset of these that are essential genes. **(B)** Correlation of protein and RNA abundance in HEK293T cells (OpenCell). For comparison purposes, RNA and protein abundances in our dataset are compared to two external references: HEK293 cell line RNASeq from the Human Protein Atlas, and the HeLa proteome published in (*7*). In both cases, our data correlates well with existing references. **(C)** Repeated from Fig. 1C. **(D)** Fluorescent detection success rates for proteins at different percentiles of abundance in the proteome. **(E)** To evaluate the influence of CRISPR editing efficiency on the ability to successfully select fluorescently tagged cells, we genotyped 432 cell lines from our library before FACS sorting (these lines were randomly selected). After FACS sorting (top 1% fluorescent cells, see Fig. S3A), all lines were imaged by fluorescence microscopy. Within this set, no fluorescence could be detected in 99 lines (23% of total). For well-expressed proteins (top $50^{th}$ percentile of abundance in the whole proteome), un-successful detection is correlated with low rates of CRISPR-mediated homologous recombination before FACS selection. A low rate of homologous recombination likely prevents the successful selection of a fluorescent pool by FACS.

**Fig. S3: Cell library characterization and quality control (related to Fig. 1). (A)** Optimization of sorting strategy. Polyclonal cell pools were sorted using gates of increasing fluorescence (left panel) and genotyped to quantify the enrichment for mNG11-inserted alleles (right panel, showing data for 6 different target genes). This informed our final sorting strategy in which the top 1% of fluorescent cells (gate I) were selected. **(B)** Genotype analysis of the polyclonal OpenCell library. A single allele is required for fluorescence, but our cell collection is enriched for homozygous insertions. In total, mNG11 insertions account for 61% (median) of alleles in a given cell pool across the full library (Boxes represent 25th, 50th, and 75th percentiles, and whiskers represent 1.5x interquartile range). The median values of mNG11 integrated alleles, *wt* alleles and other alleles are shown on the right. **(C)** Measurement of target protein abundance in final selected cell pools vs. parental cell line, by quantitative Western blotting. **(D)** Measurement of target protein abundance in final selected cell pools vs. parental cell line, by single-shot mass spectrometry. In these experiments, tagged lines are measured in a single replicate and compared to 6 replicates of non-edited control cell lines. Outliers targets are defined by an abundance that deviates by more than 2.5 standard deviations and by more than 2-fold of their abundance in the controls. The 5 outlier lines are outlined. **(E)** Distribution of Pearson correlation values measuring the overall correlation of abundances for all cellular proteins in each tagged cell line vs. median control. **(F)** For the outliers outlined in (D), correlation of abundances for all cellular proteins in the tagged cell line vs. median control. The abundance correlations for two individual control repeats are shown for reference. **(G)** Examples of overexpression artifacts. Single z-slice confocal images are shown (scale bar: 10 μm). Endogenously tagged lines and their equivalent overexpression constructs were not imaged using the same laser power, so that signal intensities are not directly comparable. Nuclei are shown as blue outlines (nuclei can be located in a different z-plane than the one shown). "Masking effects" are defined as the loss of fine localization details upon overexpression.

**Fig. S4: Interactome analysis (related to Fig. 2). (A)** Strategy for defining enrichment threshold to define interactions. Our strategy builds upon methods described by Hein et al (*7*). Here we use a quantitative approach to define enrichment thresholds dynamically for each replicate set, globally constrained by the parameter $a_{threshold}$. **(B)** To optimize parameter choice, we measured how precision (% co-localization) and recall (% CORUM coverage) of the corresponding interaction network varied with $a_{threshold}$. This informed a final value of 0.12. **(C)** Comparing interaction recall (% CORUM coverage) of OpenCell vs. other large-scale interactomes, including direct or $2^{nd}$-neighbor interactions (i.e., sharing a direct interactor in common). **(D)** Comparing interaction precision (% co-localization) of OpenCell vs. other large-scale interactomes. CORUM interactions are shown as a reference. **(E)** Direct comparison of OpenCell vs. Bioplex 3.0 on identical bait set. Both datasets use the same HEK293T cell line and share a large number (683) of baits in common. Precision and recall analysis by varying threshold for interaction detection ($a_{threshold}$ in OpenCell and *pInt* in Bioplex) is shown for the intersection set of 683 baits (dots represent values using thresholds used for final publication sets in both studies). For these set of overlapping baits, OpenCell also includes many new measured interactions for that intersection set of baits (right panel, top). The interactions unique to OpenCell have high precision values (right panel, bottom). **(F)** Compressibility analysis (*32*) of OpenCell vs. other large-scale interactomes. **(G)** Number of interactions measured in OpenCell (in the full dataset) that were also measured in Hein et al. (*7*) or BioPlex 3.0. **(H)** Distribution of GO annotation overlap between protein pairs identified in low-stoichiometry and high-stoichiometry interactions. **(I)** MCL clustering performance (F1 score) using stoichiometry-weighted or unweighted interaction graphs, derived from CORUM interactions as described in Drew et al (*90*).

**Fig. S5: Sequence analysis of orphan proteins (related to Fig. 2). (A)** Amino-acid sequence alignment between human NHSL1, NSHL2, KIAA1522 and NHS. **(B)** Correspondence of RAVE complex members in *S. cerevisiae, D. melanogaster* and *H. sapiens*. Note that in *S. cerevisiae* RAVE also includes Skp1, not depicted here.

**Fig. S6: Computer vision for automated microscopy acquisition (related to Fig. 3). (A)** To automate microscopy acquisition on 96-well plates and to limit experimental variability between imaging sessions (e.g., to limit variations in cell density) we paired an acquisition script, written in Python, with a pre-trained machine learning model to select field of views (FOVs) on-the-fly during the acquisition. A total of 25 FOVs are sampled per well in a single z-plane, and desirable FOVs are selected for further 3D confocal acquisition on the basis of a score predicted by the pre-trained model. **(B)** Microscopy automation workflow. Microscope hardware is controlled by a Python-based acquisition script via an open-source MicroManager-Python bridge (mm2python; https://github.com/czbiohub/mm2python). This approach enables us to combine custom acquisition logic with the rich ecosystem of Python-based machine-learning packages. Here, we use the scikit-image package to extract features from each FOV snapshot, then use a pre-trained random-forest regression model (scikit-learn) to predict a quality score for the FOV. This process is not computationally expensive and requires less than a second; the FOV score can therefore be used immediately to determine whether the script should acquire a z-stack or else move on to the next position. To maximize the quality of our confocal z-stacks, however, we chose to visit and score all 25 FOVs in each well, then re-visit the top-scoring FOVs for confocal z-stack acquisition.

**Fig. S7: The OpenCell image dataset (related to Fig. 3). (A)** Principle of graded localization annotation (manual annotations). **(B)** Fraction of multi-localization between cellular compartments. Complete localization annotations can be found in Table S6. **(C)** Comparison of annotated localization for proteins in OpenCell and Human Protein Atlas (HPA, version v20) datasets for which annotations are inconsistent. **(D)** Extensive literature curation allows to resolve 77% of OpenCell/HPA discrepancies (full details in Table S8). Here "direct evidence" refers to proteins for which localization has been directly measured in published studies, while "functional evidence" refers to proteins for which localization might not have been directly measured, but for which literature establishes a function that is predictive of a specific localization. For example, SCFD1 is a protein whose main known function is to regulate transport between ER and Golgi. This qualifies as "functional evidence". It is annotated as localized in the ER and Golgi in OpenCell, and in the nucleoplasm (main) and cytosol (additional) in HPA. **(E)** Comparison of annotated localization for 350 orthologous proteins in OpenCell and *S. cerevisiae* yeast (from LoQaTe (*47*)). Note that in yeast Golgi and vesicles are difficult to distinguish.

**Fig. S8: high-resolution image clusters (related to Fig. 4C). (A)** Size of clusters **C** (number of proteins in each cluster) as a function of clustering resolution. Shaded regions show standard deviations calculated from 9 separate repeat rounds of clustering, and average values are shown as a solid line. **(B), (C)** Examples of clusters of cytoplasmic **(B)** and nuclear **(C)** proteins.

**Fig. S9: Full hierarchical structure of interactome and localization datasets (related to Fig. 5).** Dendrograms represent the hierarchical relationships connecting **(A)** the full set of protein communities identified in the interactome (see Fig. 2) or **(B)** the full set of high-resolution clusters identified in the image collection (see Fig. 4C). For each dataset, an intermediate layer of hierarchy separates 18-19 modules, while an upper hierarchical layer delineates three separate branches. Modules and branches are annotated on the basis of gene ontology enrichment analysis (see Suppl. Tables 5 & 9). Right-hand panels present the topological arrangement of branches (top) and modules (bottoms) in each dataset, highlighted from the full graph of connections between interaction communities ("interactome", see Fig. 2D) or from the localization UMAP ("localization", see Fig. 4C). The color codes between interactome and localization datasets are not directly comparable (i.e. same colors are not meant to represent the same exact set of proteins). **(C)** The hierarchical structures derived from interactome (left) and localization (right) datasets are compared to the hierarchical structures derived from "scrambled" controls – that is, to the hierarchical structure that is expected by chance given the proteins present in our dataset. Controls are generated by randomly shuffling the membership of each protein between spatial clusters or interaction communities. The number of proteins in each cluster or community was preserved from the original data.

**Fig. S10: Biophysical & ontology analysis of the main branches from interactome and localization hierarchies (related to Fig.s 5 and S9). (A)** The three branches derived from the image-based hierarchy (see Fig. S9A). **(B)** Enrichment analysis of GO annotations in the hierarchical branches, testing GO term enrichment of proteins in each branch against all proteins in the interactome (Fisher's exact test, showing annotations enriched at $p < 10^{-10}$ and excluding near-synonymous annotations). **(C)** The three branches derived from the interactome hierarchy (see Fig. S9B). **(D), (E)** Enrichment analysis of GO annotations in the hierarchical branches, testing GO term enrichment of proteins in each branch against all proteins in the interactome (Fisher's exact test, showing annotations enriched at $p < 10^{-10}$ and excluding near-synonymous annotations). **(F)** Heat-map representing significance testing of biophysical properties of protein sequences in the 3 branches. P-values were obtained using Student's t-test comparing proteins belonging to a specific hierarchical branch against all proteins in the three branches. **(G)** Box plots representing the significance testing of biophysical properties described in (F). Boxes represent 25th, 50th, and 75th percentiles, and whiskers represent 1.5x inter-quartile ranges. Median is represented by a white line. ** $p < 10^{-3}$ (Student's t-test), exact p-values are shown.

**Fig. S11: Unique properties of RNA-binding proteins (RNA-BPs, related to Fig. 5). (A)** Distribution of disorder score (IUPRED2) for RNA-BPs vs. non-RNA-BPs across the whole proteome. **(B)** Distribution of protein abundance for RNA-BPs vs non-RNA-BPs across the whole proteome (left) and across OpenCell targets only (right). **(C)** Distribution of number of interactors for RNA-BPs vs non-RNA-BPs across OpenCell targets. **(D)** For each OpenCell target, the number of interactors is plotted as a function of protein abundance. The subset of targets that are RNA-BPs is highlighted on the right-hand panel. Note: for boxplots in (A), (B), (C) and (D), boxes represent 25th, 50th, and 75th percentiles, and whiskers represent 1.5x interquartile range. Median is represented by a white line. **(E)** Distribution of hydrophobicity score (gravy) across spatial clusters, comparing our data to a control in which the membership of proteins across clusters was randomized 1,000 times. Lines indicate parts of the distribution over-represented in our data vs control (**: $p < 2x10^{-3}$, Fisher's exact t-test). **(F)** Distribution of high-hydrophobicity spatial clusters (average hydrophobicity score > -0.1) in the UMAP embedding from Fig. 3D (left), and ontology enrichment analysis of proteins contained in these clusters (right). Enrichment compares to the whole set of OpenCell targets (p-value: Fisher's exact test).

**Fig. S12: Interactive data exploration at opencell.czbiohub.org. (A)** The three principal pages of the OpenCell web app. From left to right: the target page, interactor page, and gallery page. **(B)** The target page consists of three columns. The leftmost column contains the functional annotation for the target from UniProt, links to other databases, our manually-assigned localization annotations, and measures of protein expression. The middle column contains the image viewer, and the rightmost column the interaction network. **(C)** The image viewer allows the user to scroll through the confocal z-slices using a slider or to visualize the z-stack in 3D as a volume rendering. In either mode, the user can pan and zoom by clicking, dragging, and scrolling. **(D)** The interaction network can be toggled with two alternative, complementary visualizations of the target's protein interactions: a volcano plot of relative enrichment vs. p-value and a scatterplot of interaction stoichiometry vs. cellular abundance stoichiometry. In both the network view and the scatterplots, the user can click on an interactor to open the target or the interactor page for the corresponding protein.

**List of Supplementary Tables**

*Note: each Supplementary Table contains a specific "read_me" tab that describes its content in detail.*

**Table S1.**
The OpenCell library (includes target information, library design and genotype data). Related to Fig. 1.

**Table S2.**

Annotated HEK293T proteome (includes RNA and protein abundance data, biophysical properties and ontologies relevant to the analyses presented in this paper). Related to Fig. 1.

**Table S3.**
Properties of successful vs. unsuccessful edited targets. Related to Fig. 1 & S2.

**Table S4.**
The OpenCell interactome (quantitative description of interactions). Related to Fig. 2.

**Table S5.**
Clustering analysis of the interactome (analysis of MCL clustering and subsequent hierarchical analyses). Related to Figs. 2 & S9.

**Table S6.**
The OpenCell localization dataset and annotations. Related to Fig. 3.

**Table S7**
Comparison of OpenCell to Human Protein Atlas (Table S7A) or yeast (Table S7B) localization annotations. Related to Fig. 3B & S7.

**Table S8.**
Resolving discrepancies between OpenCell and Human Protein Atlas annotations by literature curation. Related to Fig. S7C.

**Table S9.**
Clustering analysis of the imaging dataset (analysis of Leiden clustering and subsequent hierarchical analyses from high-resolution clusters). Related to Figs. 4 & S9.
